# Intrahepatic crystals from elevated dietary cholesterol are sufficient to stiffen the liver

**DOI:** 10.1101/2024.12.29.630682

**Authors:** David Li, Abigail E. Loneker, Yasmine Safraou, Jamie Ford, Elaine Mihelc, Kandice R. Levental, Ilya Levental, Ingolf Sack, Paul A. Janmey, Rebecca G. Wells

## Abstract

Chronic lipid accumulation is a hallmark of metabolic dysfunction-associated steatotic liver disease (MASLD), and dyslipidemia is associated with disease progression and poorer patient outcomes. Tissue stiffening is an established fibrogenic cue, but how changes in lipid accumulation affect tissue mechanics is not fully understood. Here we show that cholesterol-containing lipid crystals stiffen steatotic liver tissue. We show that rats fed elevated dietary cholesterol develop both liquid and solid cholesterol-containing lipid crystals in the liver. While steatotic livers without crystals are softer than control livers, livers with lipid crystals show increased baseline stiffness and compression stiffening as well as increased progression to fibrosis. Lipid extracts from livers containing crystals stiffen fibrous tissue mimics, while depletion of cholesterol using methyl-ꞵ-cyclodextrin reduces both crystal abundance and tissue stiffness. Our results demonstrate in a rat model that a high cholesterol diet leads to formation of liquid and solid crystals and that cholesterol-containing crystals stiffen tissues. This work implicates lipid crystals arising from dyslipidemia as a key driver of MASLD progression. The presence of cholesterol crystals could lead to new diagnostic tools for progressive MASLD and could be a therapeutic target.

## Main

Metabolic dysfunction-associated steatotic liver disease (MASLD) refers to a spectrum of disease encompassing isolated steatosis as well as more severe pathology such as steatohepatitis (MASH), fibrosis, cirrhosis, and hepatocellular carcinoma (HCC)^1,2^. MASLD affects more than one third of the global population and is increasing in incidence, presenting substantial health and economic burdens^3^. Progression from simple steatosis to advanced disease is variable and difficult to predict, and there are few effective treatments for cirrhosis and HCC, highlighting the need to better define the underlying mechanisms of disease^2,4–7^. In particular, although both dyslipidemia and hepatic steatosis are hallmarks of MASLD^8^, the links between them and the development of advanced disease are unclear. Understanding how dyslipidemia and hepatic lipid accumulation intersect and contribute to MASLD pathogenesis may be important for predicting and treating MASLD progression^4,6^.

Changes in tissue mechanics, specifically stiffening, play a major role in the development of fibrosis in many organs^9–13^, including the liver^9,10,14^. Fat accumulation is one cause of altered tissue mechanics, but whether it causes softening or stiffening is variable and tissue specific^15–18^. While recent studies suggest that lipid droplets (LDs) soften the liver in simple steatosis^19,20^, changes in lipid droplet composition due to factors such as diet may alter the mechanics of steatosis^21^. Diet can change the hepatic lipid profile^22–24^, and previous studies in patients have shown that cholesterol and cholesterol esters (CEs) are significantly higher in MASH compared to simple steatosis^25–28^. High dietary cholesterol is associated with MASLD progression and with an increased risk of adverse outcomes in patients^29–31^, and high-fat high-cholesterol (HFHC) diets in rodent models result in marked increases in fibrosis compared to control or high-fat (HF)-only diets^22,25,32^. Birefringent lipid crystals have been observed in the livers of MASH patients^33,34^, raising the possibility that composition-dependent changes in lipid organization may be involved in MASLD pathogenesis. In atherosclerosis, solid crystals of cholesterol and CEs significantly increase the stiffness of model plaques^21^, suggesting that lipid crystals may contribute to pathogenic stiffening in MASLD.

We hypothesized that high dietary cholesterol leads to lipid crystal formation in the steatotic liver, which then promotes fibrosis by changing tissue mechanics. We used rat dietary models to investigate how cholesterol-induced changes in lipid organization affect liver mechanics. We suggest that lipid crystallization is a mechanism by which metabolic dysregulation intersects with tissue mechanics to promote fibrosis.

### Increased dietary cholesterol leads to rat liver stiffening even in the absence of fibrosis

High dietary cholesterol is associated with adverse outcomes in MASLD^29–31^, and, in rats, HFHC diets lead to marked fibrosis while HF diets do not^22,25,32^. We therefore examined the livers of HF (27.5% palm oil)- and HFHC (27.5% palm oil, 2.5% cholesterol, 2% sodium cholate)-fed rats, where the diet was switched from a control diet at 9 weeks of age, focusing in particular on liver mechanics and the characteristics of lipid droplets.

While rats on HFHC and HF diets had lower caloric intake and body weight compared to those on control diets, HFHC rat livers were lighter in color and larger and heavier than control livers (Ext Data Fig. 1a,e-h), consistent with previous reports^23^. H&E and myeloperoxidase staining of liver tissue showed significantly greater hepatic steatosis and inflammation in HFHC compared to HF rat livers (Ext Data Fig. 1b-c,i-j). Early fibrosis was seen after 9 weeks (but not 3 or 6 weeks) of HFHC feeding, while fibrosis was not present in HF or control livers (Ext Data Fig. 1d,k). Together, these results show that high dietary cholesterol exacerbates the pathological effects of liver steatosis and leads to early fibrosis.

**Fig 1:**
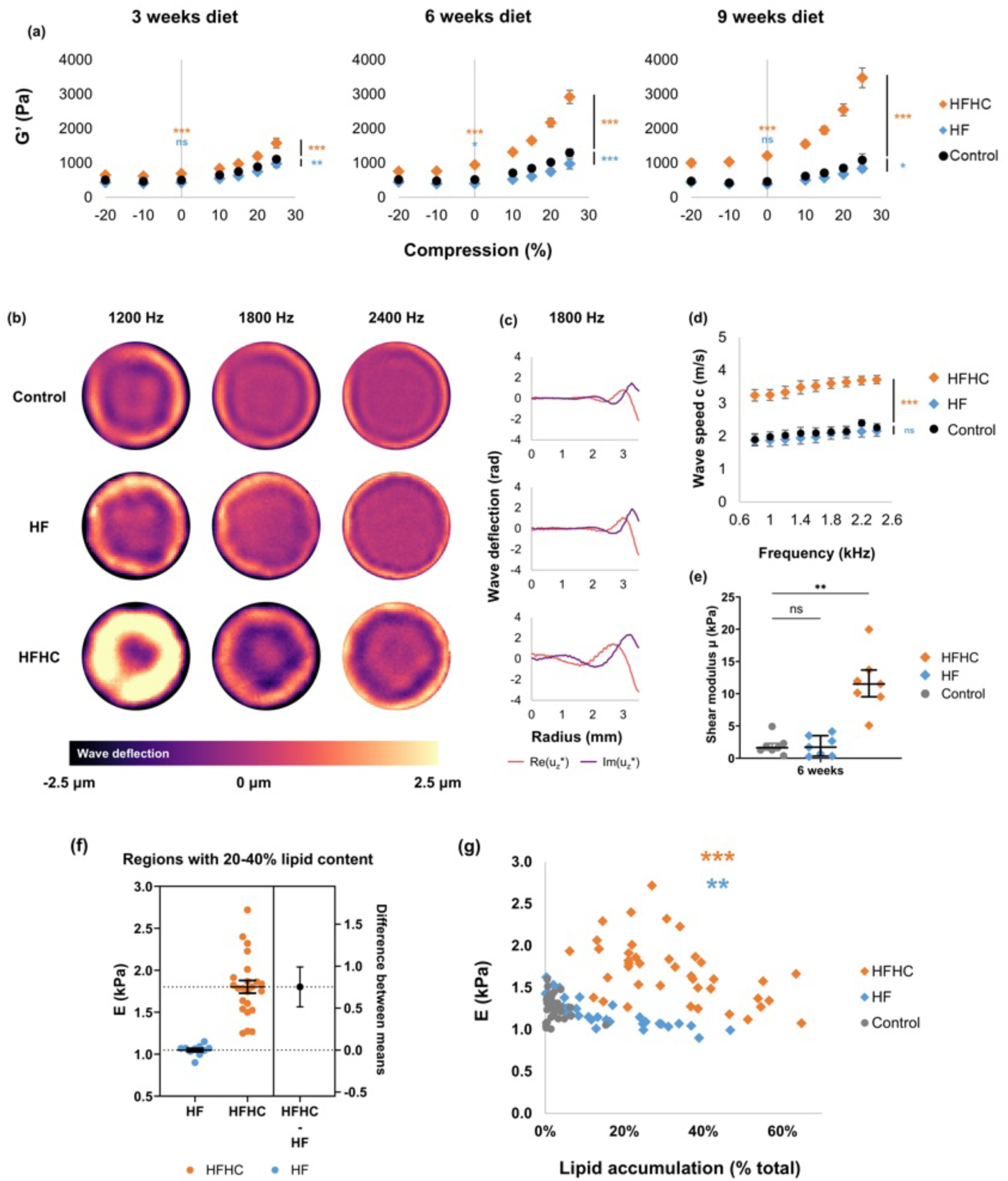
Increased dietary cholesterol results in whole liver stiffening. **(A)** Shear elastic modulus G’ of HFHC (orange), HF (blue), and control (black) livers at 3w, 6w, and 9w of diet measured by shear rheometry with variable tension and compression. HFHC livers had significantly higher shear elastic and loss moduli than control livers and greater compression stiffening, while HF livers had slightly lower shear elastic and loss moduli than control livers and less compression stiffening. n ≥ 6 total measurements from ≥ 3 animals for each condition. **(B)** Representative multifrequency wave images and **(C)** representative real (orange) and imaginary (purple) components of 2D cylinder waves at 1800 Hz from magnetic resonance elastography mapped onto a 1D axis along the sample radius of liver tissue from rats fed a control, HF, and HFHC diet for 6w, method from^38^. Shear waves penetrated deeper into HFHC liver tissue than into control or HF liver tissue at the same driving frequency. Wave images are 7 mm diameter, and color scale represents wave deflection in μm. **(D)** Wave speed c under varied driving frequencies and **(E)** shear modulus parameter μ fitted from a viscoelastic springpot model^38^ from magnetic resonance elastography of HFHC, HF, and control livers at 6w of diet. n = 7 total measurements from 3 animals for each condition. Error bars indicate standard error. * P < 0.05, ** P < 0.005, and *** P < 0.0005 for differences between curves by two-way ANOVA with Dunnett’s multiple comparisons test against control and * P < 0.05, ** P < 0.005, and *** P < 0.0005 for differences in both G’ at zero compression and shear modulus μ by one-way ANOVA with Dunnett’s multiple comparisons test against control. **(F)** Two-group estimation plot of Young’s modulus E versus diet condition (HF/HFHC) for equivalent levels of fat accumulation (between 20-40%) across 3w to 9w of diet. Error bar indicates standard error of difference between means. n = 10 and 23 for HF and HFHC, respectively. *** P < 0.001 for HF vs. HFHC local stiffnesses with equivalent levels of local fat by Student’s T-test. **(G)** Scatter plot of Young’s modulus E vs. lipid accumulation within microindented regions of HFHC (orange), HF (blue), and control (grey) livers across 3w to 9w of diet. Submillimeter regions of livers with greater local steatosis had lower local stiffness at all tested ages. n ≥ 31 measurements from ≥ 7 animals per condition. r_s_ = −0.86 and *** P < 0.001 for HF, r_s_ = −0.49 and ** P < 0.01 for HFHC by Spearman’s rank correlation.

Rheometry of rat livers showed that the HFHC diet caused tissue stiffening, with a significant increase in shear elastic modulus (G’), shear loss modulus (G”), and Young’s modulus (E) compared to control across physiological ranges of compression and tension^9,19,35^ (Fig. 1a, Ext Data Fig. 2a). Compression stiffening was particularly notable in the HFHC livers (Fig. 1a). These mechanical changes occurred even in livers after 3w and 6w of diet, when there was no detectable fibrosis. Magnetic resonance elastography provided further evidence of cholesterol-associated stiffening, with higher wave speed c and fitted shear modulus parameter μ in HFHC liver tissue (Fig. 1b-e). In contrast, the HF diet (without elevated cholesterol) led to liver softening, with a mild reduction in shear elastic, shear loss, and Young’s moduli despite elevated steatosis compared to control, in agreement with previous studies of mouse livers with simple steatosis^19,20^. The magnetic resonance elastography parameters were unchanged in HF livers compared to control, likely reflecting the higher frequency range of the elastography compared to the shear rheometry measurements (Fig. 1d, e)^36,37^.

**Fig 2:**
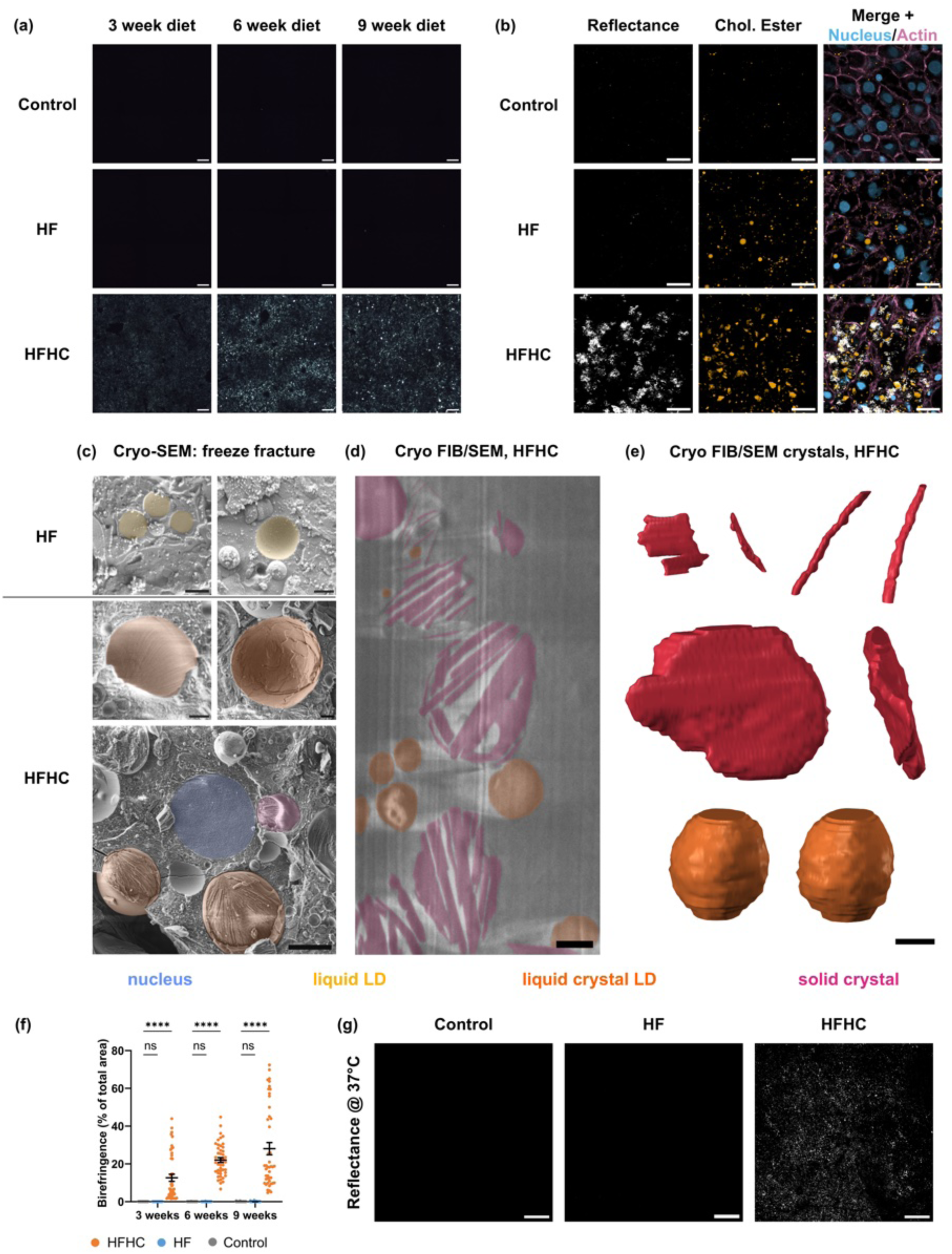
High dietary cholesterol causes localized cholesterol crystallization in the liver. **(A)** Representative frozen sections of livers from rats fed a control, HF, or HFHC diet for 3w, 6w, and 9w imaged using polarized light microscopy at 4x magnification. HFHC livers showed birefringent crystals at 3w, 6w, and 9w of diet, while no birefringence was observed for HF or control livers. n ≥ 3 animals per condition. Scale bar 200 μm. **(B)** Frozen sections of control, HF, and HFHC livers at 6w of diet imaged at 63x magnification with both confocal reflectance microscopy of reflective crystals (white) and confocal fluorescence microscopy of BODIPY-tagged cholesterol ester (orange). Slides were also imaged for actin (magenta, right) and nuclei (blue, right). Fluorescent cholesterol ester staining shows sharp, jagged crystals in HFHC liver sections and only spherical lipid droplets in HF liver sections. Regions with cholesterol ester-tagged crystals have a high reflectance signal. n = 3 livers per condition. Scale bar 20 μm. **(C)** Cryo-scanning electron microscopy (cryo-SEM) micrographs of high-pressure-frozen and freeze-fractured HF (top, blue border) and HFHC (bottom, orange border) livers at 6w of diet. HF livers contain liquid lipid droplets (liquid LDs, yellow) that either cleave cleanly or leave smooth reliefs across the fractured surface. HFHC livers contain both liquid crystals (liquid crystal LDs, orange) and solid crystals in and around LDs (red) in hepatocytes. Liquid crystal LDs of cholesterol esters can be distinguished by their characteristic onion-like internal organization with periodic 140-180 nm spacing^44^. Scale bar 1 μm for top four micrographs and 5 μm for bottom micrograph. **(D)** Representative slice of a region of high-pressure-frozen HFHC liver at 6w diet viewed by cryo-focused ion beam SEM (cryo-FIB/SEM). Solid crystals are seen within the HFHC liver, ranging from 30-200 nm in thickness and 100-5000 nm in length. Scale bar 1000 nm. **(E)** Cryo FIB/SEM segmented images of representative lipid crystals from HFHC livers at 6w diet. Each crystal reconstruction has a front and side view. Top left plate crystal is approx. 580×570×40 nm, top right rod crystal is approx. 70×70×1200 nm, middle plate crystal is approx. 2190×1580×302 nm, and bottom LD with liquid crystals is approx. 1200×1100×1200 nm. Scale bar 500 nm. **(F)** Birefringent signal as a percent of total area of livers from rats fed a control, HF, or HFHC diet for 3w, 6w, and 9w. n = 45 independent 500×500 um regions and ≥ 3 livers per condition. Error bars indicate standard error. *** P < 0.0005 and **** P < 0.0001 by two-way ANOVA with Dunnett’s multiple comparisons test against control. **(G)** Control, HF, and HFHC livers imaged using confocal reflectance microscopy at 37 °C. Livers were extracted from the rat, maintained in a 37 °C bath, and imaged <10 min from extraction. HFHC livers show strong reflectance signal from crystal organization when imaged immediately after extraction and kept at physiological temp, while no reflectance signal was observed for HF or control livers. n = 3 livers per condition. Scale bar 200 μm.

We also used a high cholesterol only (HC, 2.5% cholesterol, 2% sodium cholate, no added palm oil) diet to determine whether elevated dietary cholesterol alone was sufficient to induce stiffening. The HC diet led to tissue stiffening and fibrosis at earlier times than in animals on control diets, but to a lesser extent than for the HFHC diet (Ext Data Fig. 3). This suggested that dietary cholesterol alone can induce a MASLD-like phenotype but that the combination of elevated dietary cholesterol and fat exacerbates disease progression to a greater extent than either alone.

**Figure 3.**
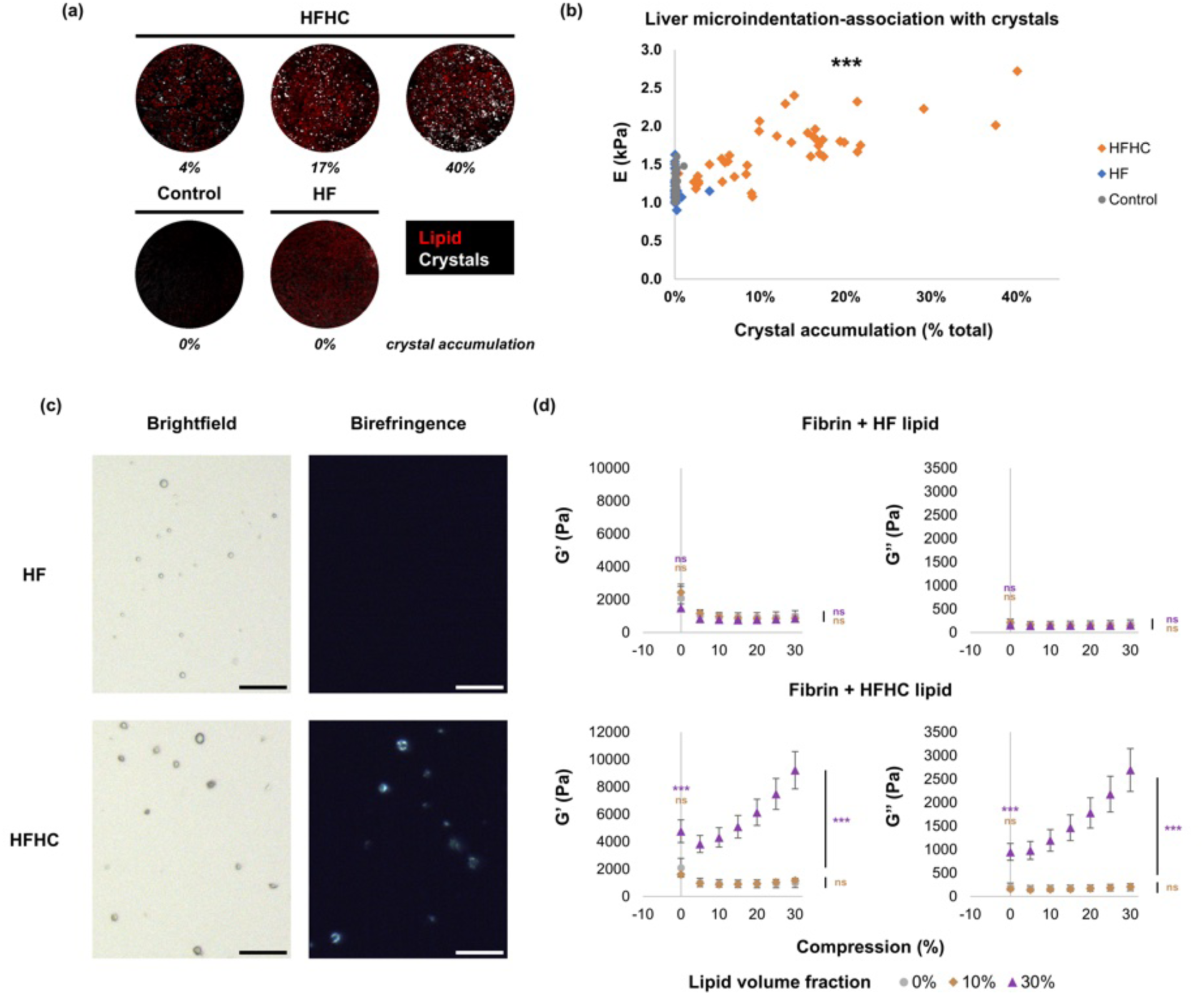
Cholesterol-containing crystals stiffen tissue and fibrin tissue mimics. **(A)** Representative confocal images of microindented regions in control, HF, and HFHC livers at 9w of diet, showing lipid accumulation using Oil Red O (red) and heterogeneous cholesterol crystal accumulation using confocal reflectance (white). Scale bar 150 μm. **(B)** Scatter plot of Young’s modulus E vs. lipid crystal accumulation within microindented regions of HFHC (orange), HF (blue), and control (grey) livers across 3w to 9w of diet. Submillimeter regions of livers with higher local crystal content had higher local stiffness at all tested ages. n ≥ 31 measurements from ≥ 7 animals per condition. r_s_ = 0.747 and *** P < 0.001 across all conditions by Spearman’s rank correlation. **(C)** Representative samples of lipids extracted from livers taken under brightfield (left) and polarized light microscopy (right) from rats fed an HF (top) and HFHC (bottom) diet for 9w. HFHC lipid crystals are highly birefringent while HF lipid droplets are not. n = 3 livers per condition. Scale bar 30 µm. **(D)** Shear storage and loss moduli under varied compression of fibrin hydrogels fabricated with 0% (grey), 10% (brown), and 30% v/v (purple) of isolated lipids from HF (top) or HFHC livers (bottom). When embedded within the model fibrin hydrogel, isolated HFHC lipid crystals at 30% v/v were sufficient to cause both significant stiffening at 0% compression and dramatically increased compression stiffening. In contrast, isolated HF lipid droplets up to 30% v/v did not alter measured mechanical properties of the composite. n = 5 hydrogels per concentration for fibrin+HFHC lipid and n ≥ 3 hydrogels per concentration for fibrin+HF lipid. Error bars indicate standard error. * P < 0.05, ** P < 0.005, and *** P < 0.0005 for differences between curves by two-way ANOVA with Dunnett’s multiple comparisons test against control and * P < 0.05, ** P < 0.005, and *** P < 0.0005 for differences at zero compression by one-way ANOVA with Dunnett’s multiple comparisons test against control.

Taken together, these results show that elevated dietary cholesterol induces stiffening of the steatotic liver before the initial development of fibrosis.

### Cholesterol-induced liver stiffening is independent of local steatosis

Despite the presence of steatosis in both HF and HFHC livers, the HFHC diet led to liver stiffening and fibrosis while the HF diet did not. HFHC diets induced significantly greater steatosis than HF diets, and we therefore tested whether the degree of steatosis determined liver stiffening. We leveraged the technique of microindentation-visualization to couple stiffness measurements to quantification of lipid accumulation by Oil Red O staining in the same submillimeter regions (Ext Data Fig. 2b)^19,39^.

In regions with similar degrees of steatosis, HFHC livers were significantly stiffer than HF livers (Fig. 1f), suggesting that cholesterol stiffens the liver independent of steatosis. Increased local lipid accumulation correlated with reduced stiffness in both HF and HFHC livers, although with a baseline of increased stiffness in the HFHC livers (Fig. 1g). These results show that steatosis alone induces softening, not stiffening, and suggests an independent role for dietary cholesterol in stiffening steatotic livers.

### Dietary cholesterol causes crystal formation in liver

Elevated cholesterol is associated with the formation of stiff crystals in atherosclerotic plaques^40,41^. We thus hypothesized that cholesterol crystals in HFHC livers were the cause of liver stiffening.

Using polarized light microscopy, we observed birefringent inclusions in HFHC and HC but not HF or control livers after 3 weeks; these increased with time on the HFHC and HC diets (Fig. 2a,f, Ext Data Fig. 4). Highly reflective inclusions in HFHC livers were located within hepatocytes, where solid crystals of cholesterol have previously been observed^33^. Furthermore, overlap between reflective inclusions and BODIPY 542/563 cholesterol ester staining suggested the presence of intracellular lipid aggregates that contain CEs, a major form of cholesterol storage (Fig. 2b). The increase in both cholesterol and CE content was demonstrated by lipidomics analysis, which showed a dramatic increase in both but not triacylglycerides in HFHC livers compared to controls (Ext Data Fig. 5). The crystallinity of these aggregates was examined by imaging using cryo-electron microscopy. Cryo-SEM of the freeze-fractured surfaces of HFHC livers showed lipid droplets with an onion-like internal organization with spacing characteristic of cholesterol ester liquid crystals^42–44^ (Fig. 2c), while cryo-FIB-SEM confirmed the presence of solid crystals associated with lipid droplets as well as freely floating in the cytoplasm (Fig. 2c-e). These crystals were highly anisotropic and mostly plate- or rod-shaped, ranging from 30-200 nm in thickness and 100-5000 nm in length, which is smaller than the size of a typical hepatocyte (13-30 μm in diameter)^45,46^. To determine whether these crystals were present at physiological temperatures, we used confocal reflection microscopy, in which the optical properties of crystals cause increased reflection of light, to visualize fresh rat livers that were freshly excised (<5 min after extraction) and kept at 37 °C in a bath^47^. The technique permitted liver imaging through the surface of an intact lobe. At 37 °C, a strong reflectance signal was observed from HFHC livers, while HF and control livers had no reflectance signal (Fig. 2g). These results show that dietary cholesterol induces the formation of both solid and liquid lipid crystals in the steatotic liver, and that these lipid crystals represent cholesterol storage.

**Figure 4.**
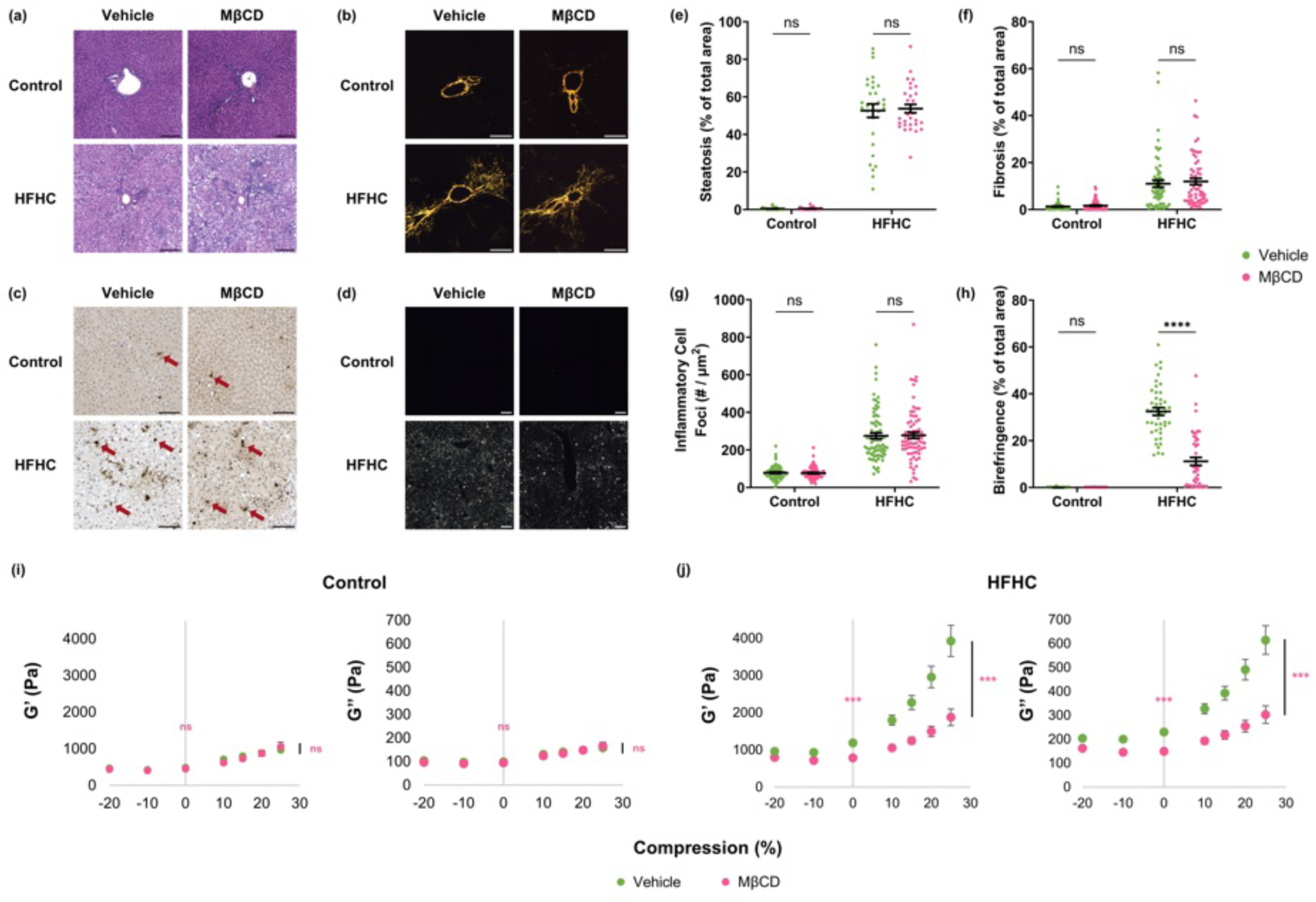
Depletion of cholesterol by MꞵCD treatment decreases tissue stiffness. **(A)** Representative H&E stains of livers from rats fed a control or HFHC diet for 9w, after in situ perfusion with either vehicle or 10 mM methyl-β-cyclodextrin (MβCD) at 37°C for 2h. Scale bar = 200 µm. **(B)** Representative picrosirius red stains of control and HFHC livers after vehicle or MβCD perfusion taken under polarized light microscopy. Scale bar = 200 µm. **(C)** Representative myeloperoxidase (MPO) immunohistochemical stains of control and HFHC livers after vehicle or MβCD perfusion. Arrows indicate representative MPO-positive inflammatory cell foci. Scale bar = 100 µm. **(D)** Representative frozen liver sections from control and HFHC livers after vehicle or MβCD perfusion imaged using polarized light microscopy. n = 3 animals per condition for (A-D). Scale bar 200 μm. **(E)** Steatosis as a percent of total area, **(F)** fibrosis as a percent of total area, **(G)** frequency of MPO-positive inflammatory cell foci in 500×500 µm regions, and **(H)** birefringence as a percent of total area of livers from rats fed a control or HFHC for 9w after vehicle or 10mM MβCD perfusion. n = 30 independent 500×500 µm regions per condition for (E), 60 independent 300×300 µm regions between adjacent central veins and portal triads per condition for (F), 75 independent 500×500 µm regions per condition for (G), and 45 independent 500×500 µm regions per condition for (H) from ≥ 3 animals per condition. * P < 0.05, ** P < 0.005, *** P < 0.0005, and **** P < 0.0001 by two-way ANOVA with Dunnett’s multiple comparisons test against control. **(I)** Shear elastic modulus G’ and **(J)** shear loss modulus G’’ of livers from rats fed a control (left) or HFHC (right) diet for 9w after in situ perfusion with either vehicle (green) or 10 mM MβCD (pink) for 2h. MβCD perfusion caused a significant reduction in both the increased liver stiffness at zero compression and increased compression stiffening caused by HFHC diet compared to vehicle perfusion. MβCD perfusion did not affect the tissue mechanics of livers from rats fed a control diet. n ≥ 8 total measurements from ≥ 3 animals for each condition. Error bars indicate standard error. * P < 0.05, ** P < 0.005, and *** P < 0.0005 for differences between curves by two-way ANOVA with Dunnett’s multiple comparisons test against control and * P < 0.05, ** P < 0.005, and *** P < 0.0005 for differences in both G’ and G’’ at zero compression by one-way ANOVA with Dunnett’s multiple comparisons test against control.

Because the presence of lipid crystals correlated with tissue stiffening in HFHC livers, we next examined whether crystals were associated with stiffening at the meso-scale. Microindentation coupled with visualization of the indented regions by confocal reflection microscopy showed that increased crystal content was strongly associated with increased stiffness in localized regions (Fig. 3a-b; compare to Fig. 1g). Thus, our results collectively demonstrate that lipid crystals are associated with liver stiffening independent of total lipid content.

### Cholesterol-containing lipid crystals directly stiffen fibrous matrices and liver tissue

We then asked whether lipid crystals directly caused tissue stiffening. Cholesterol-containing lipid crystals stiffen bulk lipid mixtures^21^; however, inflammation is another proposed cause of tissue stiffness^48,49^, so we used tissue mimics to isolate the mechanical effects of lipid crystals separate from inflammation.

We extracted lipids from HF and HFHC livers^50,51^, embedded them in 3D fibrin matrix hydrogels at physiologically-relevant concentrations^52^, and measured their effects on stiffness using shear rheometry. The fidelity of the extraction was verified by polarized light microscopy, which showed that lipids extracted from HFHC but not HF livers were highly birefringent (Fig. 3c). When the extracted lipids were embedded in the tissue mimics at 30% (v/v), mimics containing HFHC lipids had increased shear elastic and shear loss moduli, with marked compression stiffening. HF lipids had no effect on compression stiffening until added at 60% v/v (Fig. 3d, Ext Data Fig. 6), demonstrating that crystalline HFHC lipids exert a direct stiffening effect on their surrounding matrix.

We then perfused HFHC livers *in situ* for two hours with methyl-ꞵ-cyclodextrin (MꞵCD), which depletes cholesterol from both cells *in vitro* and tissues *in vivo*^53–56^. This treatment significantly decreased the birefringent signal in HFHC livers, showing that depletion of cholesterol with MꞵCD led to a significant reduction in lipid crystal content (Fig. 4d, h). Perfusion led to a marked decrease in shear elastic and shear loss moduli in HFHC livers compared to vehicle perfusion, which did not occur in control livers (Fig. 4i-j). There was no change in steatosis, fibrosis, cell shape, or inflammation, suggesting that the changes in mechanics were independent of these factors (Fig. 4a-c, e-g). The lack of change in moduli of the control livers after perfusion suggests that changes to cell membranes are not the cause of the effects we observe in the HFHC livers with perfusion.

Together, these results show that lipid crystals directly stiffen the surrounding matrix in vitro and in vivo.

## Discussion

We show here that a high cholesterol diet results in the formation of solid and liquid cholesterol-containing lipid crystals in the liver, causing liver stiffening. Our results demonstrate that lipid crystals contribute directly to soft tissue stiffening, representing a mechanical mechanism by which hyperlipidemia could drive fibrosis and MASLD progression.

Despite similar calorie intake and weight gain compared to rats on a control diet or HF diet, rats on HFHC and HC diets developed fibrosis. The time of progression to fibrosis was different between HFHC and HC diets, suggesting that lipid metabolism and composition could influence the dynamics and extent of crystal formation. Our models are similar to dietary models in mice presented as models of “lean” MASLD; cholesterol accumulation, lipid crystallization, and the resulting increased tissue stiffness may thus be a mechanism of MASLD progression in non-obese populations^23^.

The dramatic increases in cholesterol and CEs observed in HFHC livers in our lipidomics analysis are reflected by their presence in HFHC lipid crystals. A shift towards cholesterol and CEs causes ordering of lipid droplets into liquid crystals^43^, and solid lipid crystals have been reported to form on the surface of the droplets before being shed into the cytoplasm^40^. Although our cryo-FIB-SEM studies were not designed to detect extracellular cholesterol crystals, these are likely to be present as the result of the death of crystal-laden hepatocytes^33^. Microindentation-visualization showed that lipid crystals were associated with both the localization and degree of tissue stiffening whereas total neutral fat accumulation was associated with tissue softening, suggesting that crystallization changes the effect of lipid accumulation to cause tissue stiffening.

The use of fibrin gels as tissue mimics confirmed that lipid crystals in a fibrous matrix increased shear elastic and Young’s moduli both with and without compression, consistent with observations in liver. Interestingly, non-crystalline lipid droplets also increased compression stiffening, but required a much higher volume percentage of inclusions. The dramatic stiffening effect of lipid crystals at lower lipid content are likely explained by the rod- and plate-like shapes of solid crystals. Anisotropic shapes exert a greater radius of influence and excluded volume than equi-volume spherical inclusions, and can align, which would be expected to alter their mechanical influence at a larger scale. Anisotropic lipid crystals may therefore have different effects on tissue and matrix mechanics than equivalent amounts of lipid in amorphous lipid droplets^57^. Shape anisotropy also potentially decreases the percolation threshold^58,59^, explaining why crystals mediate an increase in shear and elastic moduli without compression. Finally, previous studies show that inert spherical particles embedded in a fibrous matrix lead to compression stiffening only at volume fractions above 50%, whereas spheres that bind the matrix do so at <30%, representing another potential cause for stiffening from liquid cholesterol-containing lipid crystals ^52^. Together, our results suggest that while large volumes of LDs in isolated steatosis could stiffen the liver, cholesterol-containing crystals substantially reduce the degree of steatosis required for pro-fibrotic stiffening.

The crystals we observed resemble the birefringent crystals previously noted in biopsies of human MASH patients^33,34^, highlighting the potential clinical relevance of our results. The progression from isolated steatosis to MASH and more severe disease in MASLD is highly variable^6,60^. Our finding that crystals stiffen tissues suggests that lipid crystals represent a risk factor in MASLD, and could serve as a new diagnostic tool for assessing the risk of progression. The structure and optical properties of lipid crystals may lend themselves to minimally invasive or non-invasive diagnostics^47^, such as using MRI to infer the degree of crystal formation. There are also therapeutic implications, such that low or average cholesterol diets as well as drugs that specifically target cholesterol crystals could prevent or reverse MASLD progression. While some β-cyclodextrins exhibit side effects such as ototoxicity^61^, making them unlikely (without modifications) to be useful therapeutics in MASLD, other approaches to sequester cholesterol or disrupt lipid crystallization could reduce the abundance of crystals in the liver and thereby decrease profibrotic tissue stiffening.

Tissue stiffening is a known risk factor for hepatocellular carcinoma (HCC) and other cancers^62,63^ and cholesterol-containing lipid crystals might therefore also influence the progression of MASLD to HCC^62^. Lipid crystals as a mechanism of pro-oncogenic stiffening may also be relevant to other cancers in which lipid crystals are observed, such as renal cell carcinoma, squamous cell lung cancer, and prostate cancer^64^. In addition to cancer, lipid crystals are present in and associated with a wide range of other pathologies, including diabetic retinopathy^65^ and preeclampsia^66^.

In addition to directly increasing tissue stiffness, lipid crystals could have other effects on the liver. Lipid crystals have a proinflammatory effect in other systems such as atherosclerosis^47^, and are associated with hepatic inflammation in both MASLD and in our rodent models, suggesting a second hit in the form of inflammation^33^. Additionally, we previously demonstrated cytoskeletal disruption, nuclear deformation and hepatocyte dysfunction in cells with crystal-free LDs^67^, and cholesterol-containing crystals may cause similar cytoplasmic and nuclear damage as well as physical disruption of lipid membranes such as the cell membrane or nuclear membrane^68^.

In conclusion, we demonstrate in a rat model that high dietary cholesterol leads to steatosis, fibrosis, and intra-hepatocyte crystal formation, with crystal-mediated stiffening of the liver. Cholesterol accumulation and lipid crystallization in the liver may represent a new mechanism of disease as well as a diagnostic and therapeutic target.

## Extended Data

**Extended Data Figure 1:**
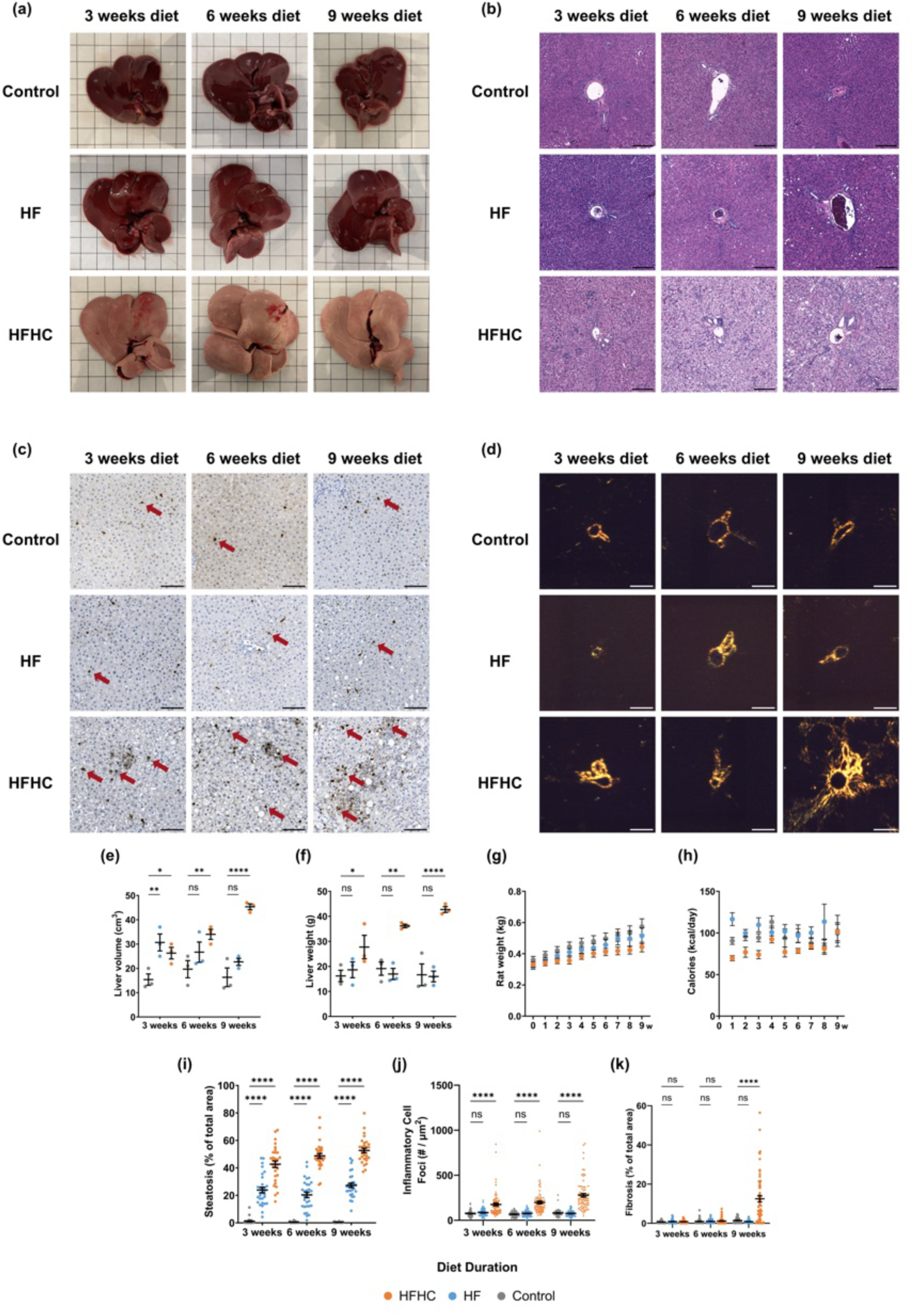
Increased cholesterol intake leads to increased steatosis and fibrosis in rats. **(A)** Representative gross images of livers from rats fed a control, high-fat (HF), or high-fat-high-cholesterol (HFHC) diet, starting at 9w of age, for 3w, 6w, and 9w. Boxes in grid are 1×1 cm. **(B)** Representative H&E stains of control, HF, and HFHC livers. Scale bars = 200 µm. **(C)** Representative myeloperoxidase (MPO) immunohistochemical stains of control, HF, and HFHC livers. Arrows indicate representative MPO-positive inflammatory cell foci. Scale bars = 100 µm. **(D)** Representative picrosirius red stains of control, HF, and HFHC livers imaged using polarized light microscopy. Scale bars = 200 µm. **(E)** Gross liver volume, **(F)** gross liver weight, **(G)** rat weight, **(H)** caloric intake, **(I)** steatosis as a percent of total area, **(J)** frequency of MPO-positive inflammatory cell foci in 500×500 µm regions, and **(K)** fibrosis as a percent of total area of livers from rats fed a control, HF, or HFHC diet for 3w, 6w, and 9w. n = 3 livers per condition for (A-F and I-K), 6-26 rats per condition for (G-H), 30 and 75 independent 500×500 um regions per condition for (I-J), and 60 independent 300×300 um regions between adjacent central veins and portal triads per condition for (K). Error bars indicate standard error. * P < 0.05, ** P < 0.005, *** P < 0.0005, and **** P < 0.0001 by two-way ANOVA with Dunnett’s multiple comparisons test against control.

**Extended Data Figure 2:**
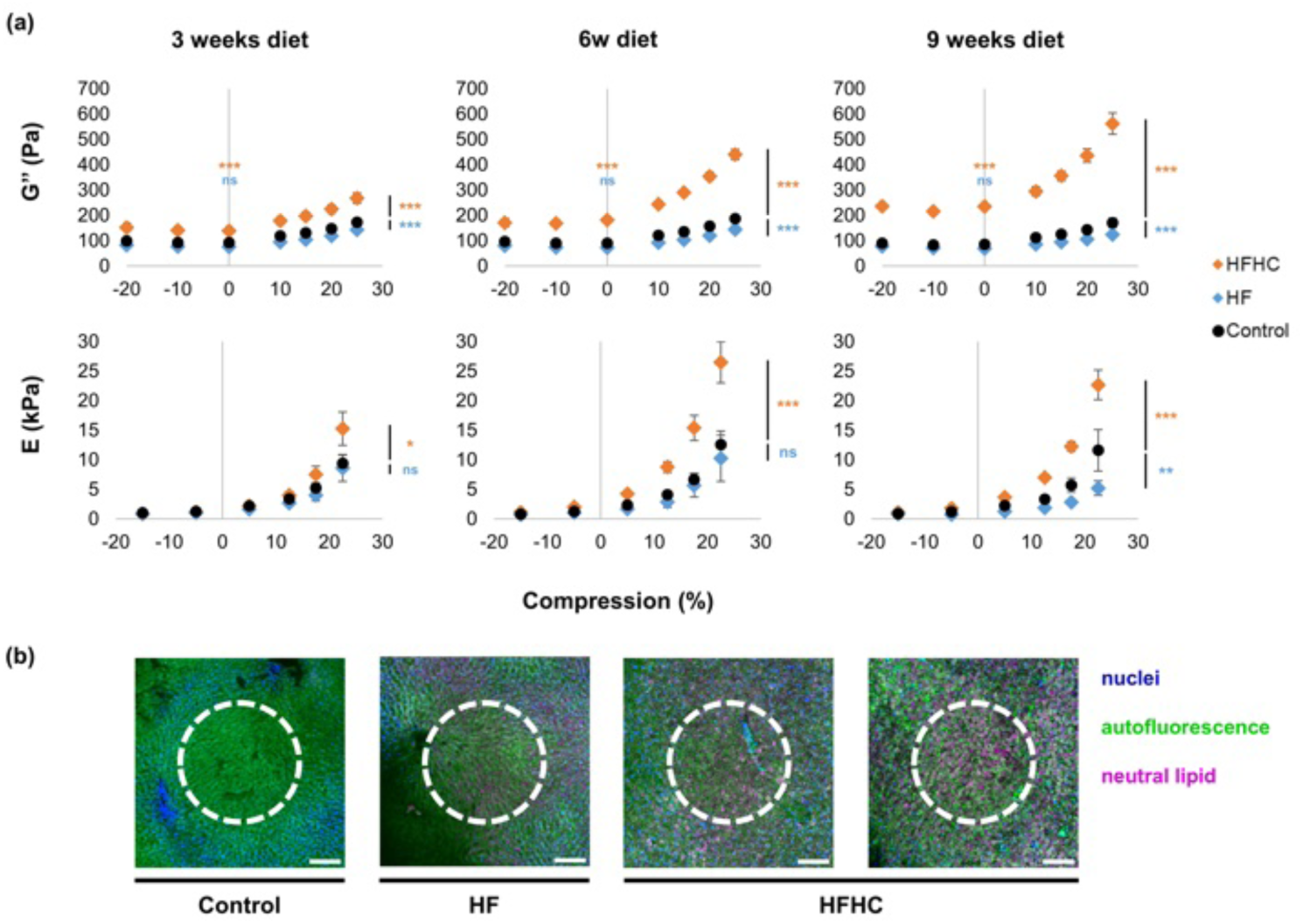
Dietary cholesterol is associated with increased Young’s modulus even without fibrosis. **(A)** Shear loss modulus G’’ and Young’s modulus E of HFHC (orange), HF (blue), and control (black) livers at 3w, 6w, and 9w of diet under varied tension and compression. HFHC livers had significantly higher Young’s moduli than control livers and greater compression stiffening, while HF livers had slightly lower Young’s moduli than control livers and less compression stiffening. n ≥ 6 total measurements from ≥ 3 animals for each condition. Error bars indicate standard error. * P < 0.05, ** P < 0.005, and *** P < 0.0005 for differences between curves by two-way ANOVA. **(B)** Representative fluorescent images of indented control, HF, and HFHC livers at 3w of diet with equivalent fat accumulation within the indented region, showing DAPI (blue) demarcating the microindented regions (white dashed line), autofluorescence (green), and heterogeneous lipid accumulation using Oil Red O (magenta). Scale bar 150 μm.

**Extended Data Figure 3:**
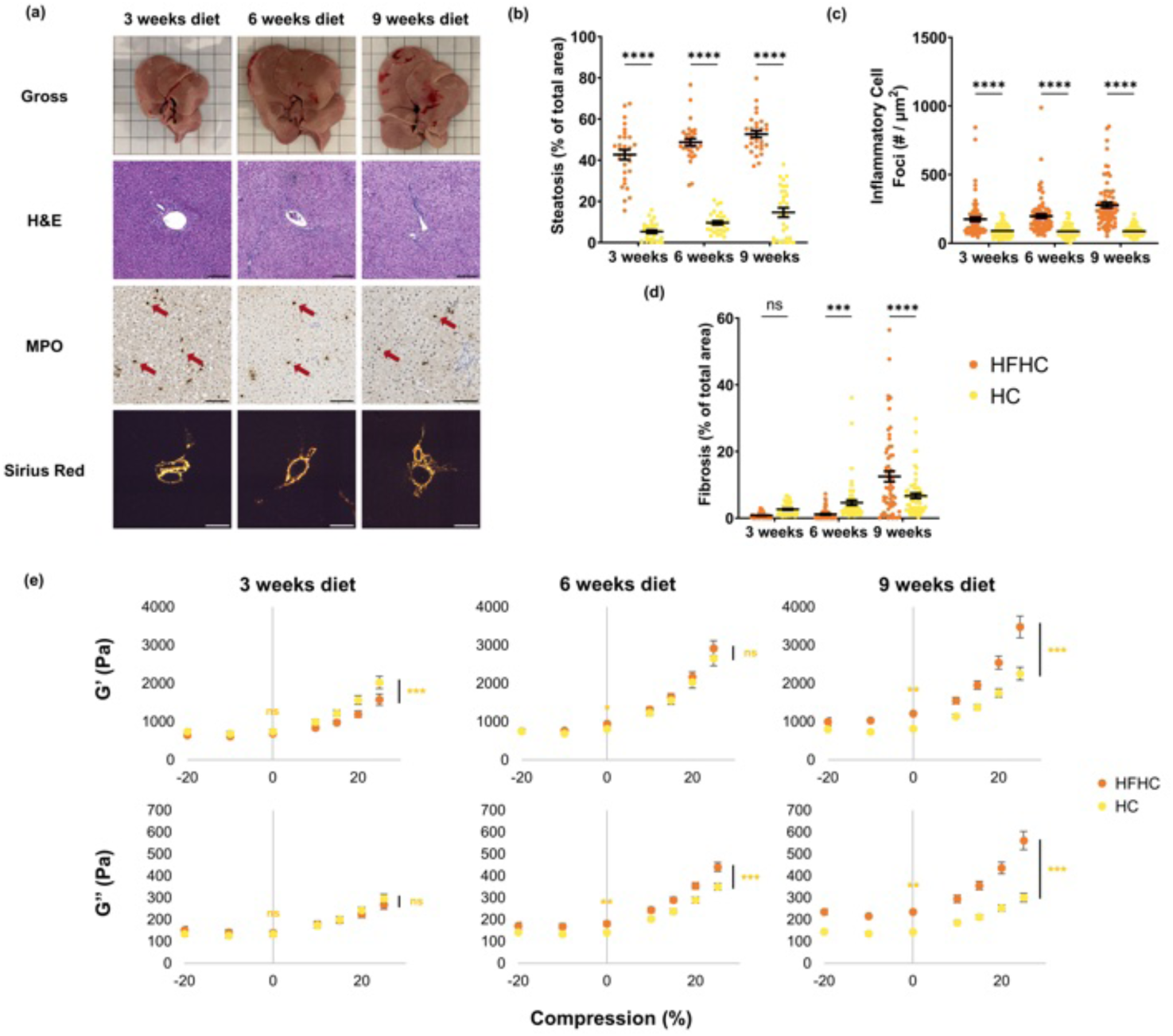
High dietary cholesterol induces liver steatosis, fibrosis, and tissue stiffening. **(A)** Representative gross images, H&E stains, myeloperoxidase IHC stains and picrosirius red stains of livers from rats fed a high-cholesterol-only (HC) diet starting at 9w of age for 3w, 6w, and 9w. Picrosirius red stains were taken under polarized light microscopy. Boxes for gross images are 1×1 cm, scale bars are 100 µm for MPO IHC stain images and 200 µm for all others. **(B)** Steatosis as a percent of total area, **(C)** frequency of MPO-positive inflammatory cell foci in 500×500 µm regions, and **(D)** fibrosis as a percent of total area of livers from rats fed a HFHC or HC diet for 3w, 6w, and 9w. Arrows indicate representative MPO-positive inflammatory cell foci. n = 30 independent 500×500 um regions per condition for (B), 45 independent 500×500 μm regions per condition for (C), 75 independent 500×500 μm regions per condition for (D), and 60 independent 300×300 μm regions between adjacent central veins and portal triads per condition for (E) with three animals per condition. Error bars indicate standard error. * P < 0.05, ** P < 0.005, *** P < 0.0005, and **** P < 0.0001 by two-way ANOVA with Dunnett’s multiple comparisons test against HFHC diet as comparison control.**(E)** Shear elastic modulus G’ and shear loss modulus G’’ of HFHC and HC livers after 3w, 6w, and 9w diet under varied tension and compression. HC livers were somewhat stiffer than HFHC livers at 3w, but had comparable stiffness to HFHC livers at 6w and were less stiff than HFHC livers at 9w. n ≥ 8 total measurements from ≥ 3 animals for each condition. Error bars indicate standard error. * P < 0.05, ** P < 0.005, and *** P < 0.0005 for differences between curves by two-way ANOVA with Dunnett’s multiple comparisons test against control and * P < 0.05, ** P < 0.005, and *** P < 0.0005 for differences in both G’ and G’’ at zero compression by one-way ANOVA with Dunnett’s multiple comparisons test against HFHC diet as comparison control.

**Extended Data Figure 4.**
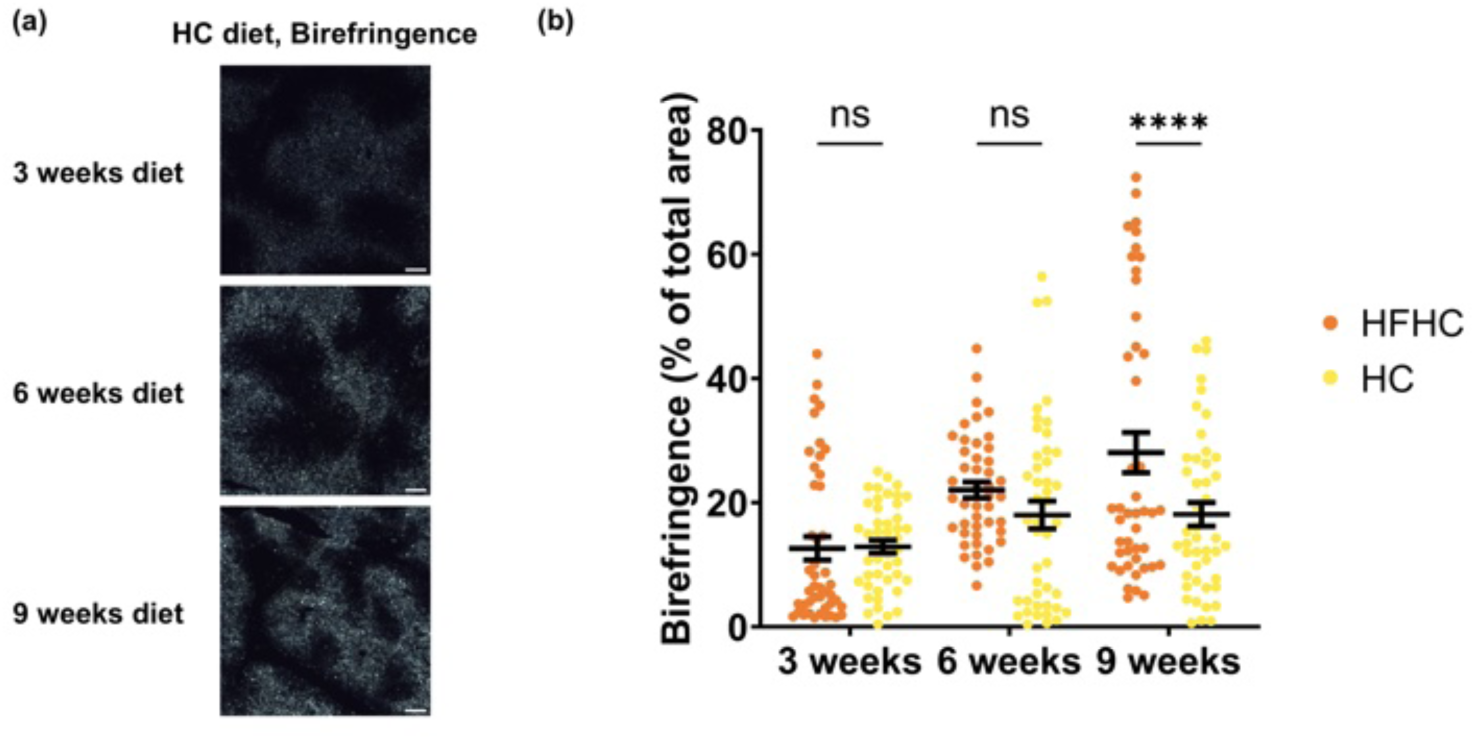
A high cholesterol diet induces lower hepatic lipid crystal accumulation than a HFHC diet. **(A)** Representative frozen sections of livers taken under polarized light microscopy from rats fed a high-cholesterol-only (HC) diet starting at 9w of age for 3w, 6w, and 9w. Scale bars are 200 µm. **(B)** Birefringence as a percent of total area of livers from rats fed a HFHC or HC diet for 3w, 6w, and 9w. 45 independent 500×500 um regions per condition with three animals per condition. Error bars indicate standard error. * P < 0.05, ** P < 0.005, *** P < 0.0005, and **** P < 0.0001 by two-way ANOVA with Dunnett’s multiple comparisons test against HFHC diet as comparison control.

**Extended Data Figure 5.**
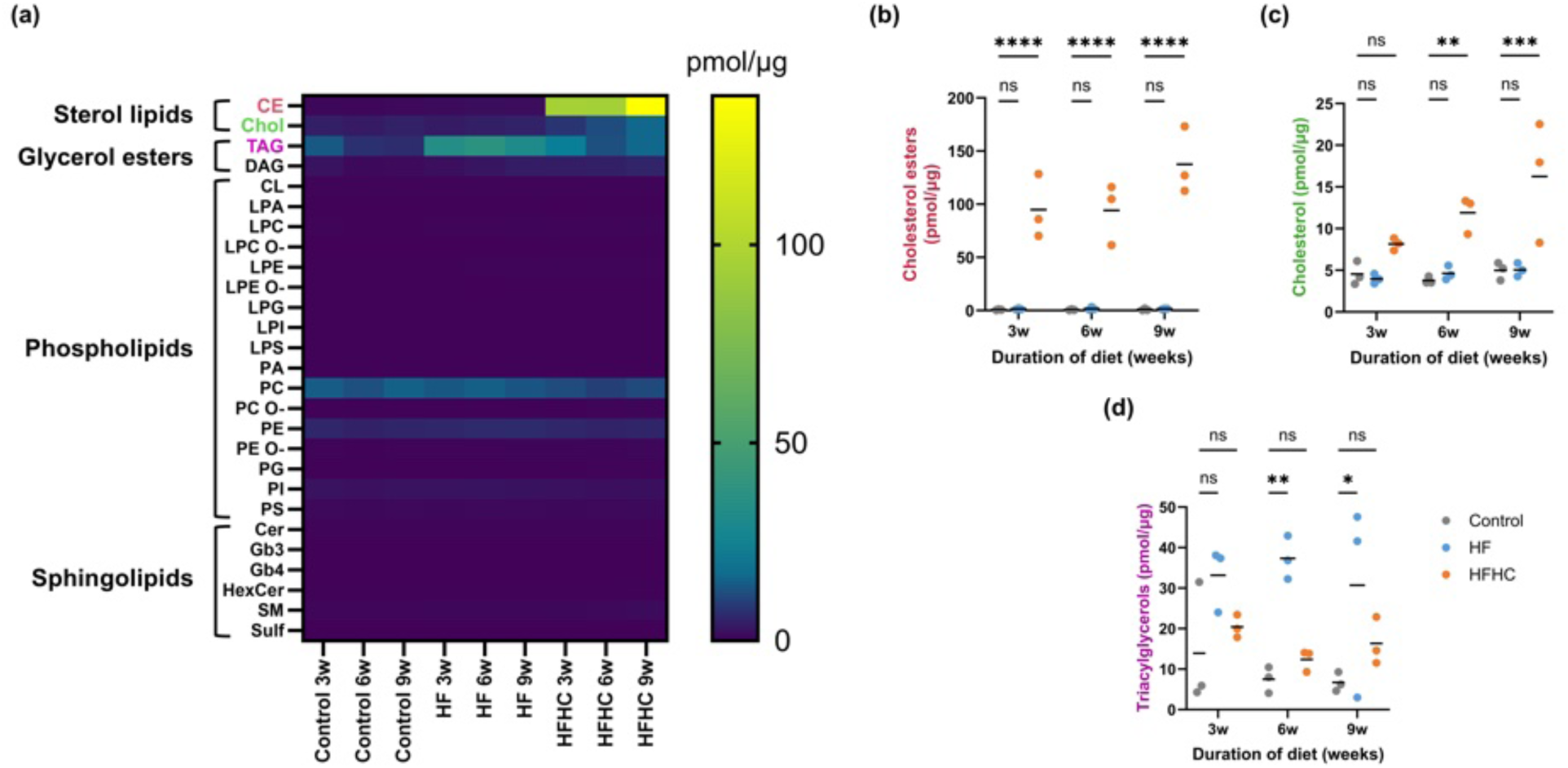
Cholesterol and cholesterol esters accumulate in the livers of rats fed a HFHC diet. **(A)** Heat map of lipid species from lipidomic analysis of the livers of rats fed a control, HF, or HFHC diet for 3w, 6w, and 9w in pmol/μg wet weight tissue. Abbreviations: CE – cholesterol esters, CL – cardiolipin, Cer – ceramide, Chol – cholesterol, DAG – diacylglycerol, HexCer – hexosylceramide, LPA – lyso-phosphatidate, LPC – lyso-phosphatidylcholine, LPE – lyso-phosphatidylethanolamine, LPG – lyso-phosphatidylglycerol, LPI – lyso-phosphatidylinositol, LPS – lyso-phosphatidylserine, PA – phosphatidate, PC – phosphatidylcholine, PE – phosphatidylethanolamine, PG – phosphatidylglycerol, PI – phosphatidylinositol, PS – phosphatidylserine, SM – sphingomyelin, Sulf – sulfatide, TAG – triacylglycerol, O- – (-ether). n = 3 livers per condition. **(B)** Cholesterol ester, **(C)** Cholesterol, and **(D)** Triacylglycerol levels of livers from rats fed a control, HF, or HFHC diet for 3w, 6w, and 9w. n = 3 livers per condition. Line indicates mean. * P < 0.05, ** P < 0.005, *** P < 0.0005, and **** P < 0.0001 by two-way ANOVA with Dunnett’s multiple comparisons test against control.

**Extended Data Figure 6.**
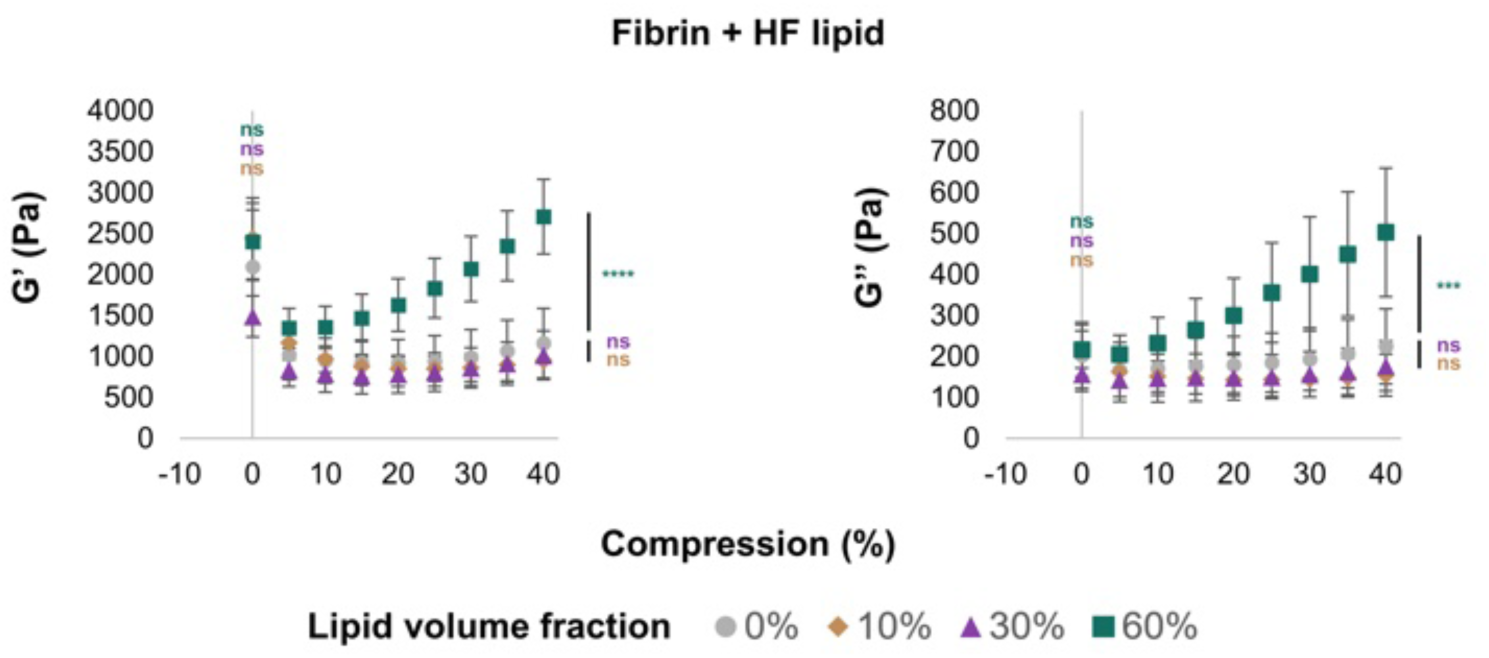
HF-associated lipid droplets in fibrous matrices have compression stiffening at high volume fractions. Shear storage G’ and shear loss G’’ moduli under variable compression of fibrin hydrogels embedded with 0% (grey), 10% (brown), 30% v/v (purple), and 60% v/v (green) of isolated lipids from HF livers. Isolated HF lipid droplets caused compression stiffening of the hydrogel only at 60% v/v. n ≥ 3 hydrogels per concentration. * P < 0.05, ** P < 0.005, and *** P < 0.0005 for differences between curves by two-way ANOVA with Dunnett’s multiple comparisons test against control and * P < 0.05, ** P < 0.005, and *** P < 0.0005 for differences at zero compression by one-way ANOVA with Dunnett’s multiple comparisons test against control.

## Materials and Methods

### Animal studies

All animal work was carried out in strict accordance with the recommendations in the Guide for the Care and Use of Laboratory Animals of the National Institutes of Health. Animal protocols were approved by the Institutional Animal Care and Use Committee of the University of Pennsylvania (protocol #804031). Sprague Dawley rats were obtained from Charles River Laboratories (strain #001) and were housed in a temperature-controlled environment with appropriate enrichment, 12:12 h light/dark cycles, and ad libitum access to water and food. Additional cholesterol and fat were added to a standard chow diet using previously published ratios to generate steatosis, steatohepatitis, and liver fibrosis^32,69^. Nine-week old rats of mixed genders were randomly assigned into cohorts and fed control chow (LabDiet, 5001), a high fat-only diet (HF) (Animal Specialties and Provisions, 27.5% palm oil, 5ZFD), a high cholesterol-only diet (HC) (Animal Specialties and Provisions, 2.5% cholesterol, 2% sodium cholate, 5ZP3), or a high-fat-and-high-cholesterol diet (HFHC) (Animal Specialties and Provisions, 27.5% palm oil, 2.5% cholesterol, 2% sodium cholate, 5Z6Z). Energy intake and body weight were recorded weekly, and cage positions were shuffled randomly every week. Euthanasia at 3, 6, and 9 weeks of diet treatment was carried out by CO_2_ inhalation followed by exsanguination. Whole livers were harvested and stored in PBS at 4 °C for up to 5 h until analysis; rodent livers are adequately preserved under these conditions and rheological properties are maintained^35,70^. Gross images of the livers were taken at the time of euthanasia. No animals were excluded during the study.

### Histology and immunohistochemistry

Histology, immunohistochemistry, and quantification of steatosis, inflammation, and fibrosis were performed as described previously^19^. Paraffin-embedded (FFPE) liver sections were stained with either hematoxylin and eosin or picrosirius red (Poly Scientific R&D Corp, S2365). Immunohistochemical staining was performed as described previously^71^ with rabbit monoclonal anti-myeloperoxidase (MPO) antibody (1:1000, Abcam, Cat# ab208670, RRID: AB_2864724) as the primary antibody^72^. Liver samples were also snap frozen in OCT and sectioned with a cryostat before staining with 10 μM BODIPY 493/503 (Thermo Fisher, D3922) and 1 μg/mL DAPI (Thermo Fisher, D1306) for 30 min at room temperature. The degree of steatosis was quantified by measuring the area of BODIPY staining in multiple 500 × 500 μm regions randomly selected throughout the liver. The degree of inflammation was quantified as the number of MPO-positive inflammatory cell foci in 500 × 500 μm regions randomly taken throughout the liver, while the degree of fibrosis was quantified by measuring the area of picrosirius red signal under crossed circular polarizers in multiple 300 × 300 μm regions between adjacent central veins and portal triads. Sample sizes were determined based on previous literature^19^, and analysis was performed blinded. Picrosirius red signal under polarized light microscopy is an established technique to maximize the sensitivity of fibrosis detection in FFPE tissue slides^73^.

Frozen HFHC liver sections were also permeabilized with 0.2% v/v Triton-X and stained with 165 nM Alexa Fluor 647 Phalloidin (Thermo Fisher, A22287), 3.8 μM CholEsteryl BODIPY 542/563 C11 (Thermo Fisher, C12680), and 1 μg/mL DAPI for 1 hr at 37°C for identification and visualization of cholesterol ester crystals^64,65,74^.

### Shear rheometry

Shear rheometry was performed as described previously^19,35^. Briefly, liver samples were prepared using an 8 mm punch (Alabama R&D, MP0144) with all punches taken in the same orientation from the same lobe. The height of the samples ranged from 4.7 to 7.7 mm in the uncompressed state. Samples were kept hydrated with PBS during all experiments. Parallel plate shear rheometry was carried out on a Kinexus PRO rheometer (Malvern Instruments, Kinexus series) at room temperature. Samples were attached to rheometer platforms with fibrin glue by applying 5 μL each of 20 mg/mL bovine plasma fibrinogen (Sigma-Aldrich, 341573) and 100 U/mL bovine thrombin (Sigma-Aldrich, T4648) to both the top and bottom surfaces of the sample. The upper platen was quickly lowered until 0.02 N of nominal initial force was applied to ensure adhesive contact of the sample with the metal surfaces of the rheometer, and the sample was allowed to sit for 10 min to allow the fibrin glue to polymerize fully before performing measurements. The fibrin glue does not affect the mechanics of the system^35^. Mechanics were measured with a dynamic time sweep test (2% constant strain, oscillation frequency 1 rad/s, measurements taken for 120 s). These measurements were carried out first uncompressed, then with increasing uniaxial tension (10% and 20%), then uncompressed, then with increasing uniaxial compression (10, 15, 20, and 25%). Tension and compression were applied by changing the gap between the platform and upper platen of the rheometer. Previous studies show that changes in liver mechanics from these levels of tension and compression are reversible^35^. The following correction was applied to account for the change in cross-sectional area during testing under the assumption that total tissue volume is conserved, where G′ is the storage modulus and λ is axial strain:

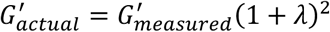

A similar correction was applied to the loss modulus G′′ and compressive Young’s modulus E.

### Magnetic resonance elastography

Magnetic resonance elastography of liver cores was performed as described previously^38,75,76^. Briefly, a 0.5-T compact tabletop MRI scanner (Pure Devices GmbH) with a 10-mm bore and 0.5-T permanent magnet was customized by addition of an external gradient amplifier (Pure Devices GmBH, DC 600) and an integrated MRI system-controlled piezoelectric driver (Piezosystem Jena). The piezoelectric driver was fed with a sinusoidal alternating current of maximum 90 V (depending on the desired vibration amplitude) and frequencies between f = 800 and 2400 Hz from the gradient amplifier. A glass tube with inner diameter 7 mm was mounted onto the piezoelectric driver and shear waves were introduced into tissue samples from the cylinder walls by producing concentric cylinder waves emanating from the outer sample boundaries toward the center of the tissue. Liver tissue was imaged using the following acquisition parameters: slice thickness = 3 mm, matrix size = 64×64, field of view = 9.6 mm, resolution = 0.15 x 0.15 x 3 mm^3^, and driving frequencies between 800 and 2400 Hz, with a 200 Hz step.

For each driving frequency f, the acquired phase data were unwrapped and Fourier-transformed in time to extract complex-valued wave images. These wave images were fitted by an analytical solution based on Bessel functions prescribed in a z-infinite cylinder. The complex-valued wave number was translated into two real-valued quantities related to stiffness (shear wave speed c) and viscous attenuation (shear wave penetration rate a), both in m/s:

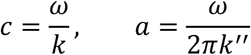

where ω is the angular driving frequency equal to 2πf and k*=k′+ik′′ is the complex-valued wave number. The viscoelastic springpot model, which is a two parameter fractional element, was fitted on shear wave speed as a function of frequency, yielding a frequency-independent shear modulus μ in kPa and dimensionless power law exponent ɑ:

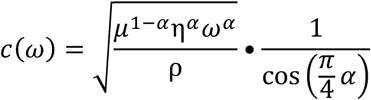

assuming specific viscosity η = 1 Pa*s and tissue density ρ= 1 kg/L.

### Microindentation, demarcation, and visualization

Microindentation was performed as described previously to permit the demarcation and visualization of mesoscale features within the microindented regions of the tissue^19,39^. Multiple measurements were obtained per liver sample using this approach. After microindentation and demarcation, liver samples were fixed in 10% neutral buffered formalin for 12 h at 4 °C before storage in PBS until staining and imaging for lipid and crystal accumulation. The investigator was blinded to the identity of the samples during measurements. Liver samples were stained for lipid using an adaptation of whole mount Oil red O staining^77^ at room temperature on a shaker. Crystal accumulation was visualized using confocal reflection microscopy, as below.

### Confocal reflection microscopy

Confocal reflection microscopy was performed in tandem with confocal fluorescence microscopy as described previously^47^. Reflection was captured by placement of the detector channel directly over the wavelength of the selected laser channel. The laser channel used for reflection was chosen to prevent spectral overlap with the fluorescent channels used for standard confocal fluorescence.

To obtain warm, fresh livers for visualization of crystals, rats were anesthetized with 100 mg/kg body weight pentobarbital sodium (Sagent Pharmaceuticals, 25051-676-20) and monitored until the rat was confirmed to have reached a surgical plane of anesthesia. The abdominal cavity was then opened to expose the liver under a heat lamp adjusted to keep the organs at 37 °C. While the rat was euthanized via exsanguination, the superior right lateral lobe was removed and transported into the confocal microscope in a 50 mL conical of PBS within < 5 min while maintained at 37 °C. The lobe was then laid onto a glass coverslip and moist Kimwipes were placed on top to maintain humidity. The entire microscope and microscope stage was maintained at 37 °C throughout the duration of observation. Crystals within the liver were imaged directly through the intact Glisson’s capsule/liver lobe surface without further processing or cutting.

### Cryo-EM sample preparation

Livers were fixed for cryo-EM similar to previously published protocols^40,78^. Briefly, rats were anesthetized and livers were flushed with 1,000 USP units/mL heparin (Fresnius Kabi, 504105) and briefly with HBSS w/ Ca,Mg (Thermo Fisher, 14025) before perfusion with 1.5% glutaraldehyde (Millipore Sigma, G5882) and 1% sucrose (Millipore-Sigma, S0389) in 0.067 M cacodylate buffer (Electron Microscopy Sciences, 11653) at 15 mL/min for 7 min^78^. All perfusion solutions were maintained at 37 °C in a warm water bath for the duration of the perfusion. Rat liver temperature was regularly measured using an infrared thermometer and maintained at 37 °C for the duration of the perfusion using an infrared heat lamp. After perfusion, the rat liver was removed, and samples prepared for histology as above.

Liver tissue cores were obtained using an 8 mm punch, the liver capsule was removed, and cores were sliced to a thickness of ∼200 μm using a Brendel/Vitron tissue slicer (Vitron). Fixed liver slices were incubated in 0.22 μm-filtered 10% dextran (Sigma-Aldrich, 31389) in PBS for 70 min prior to high pressure freezing to allow for cryoprotectant infiltration^79^. Afterwards, 2 mm diameter tissue pieces were extracted from the liver slices, sandwiched between brass planchettes, and subjected to high pressure freezing using a Bal-Tec ABA HPM 010 high pressure freezer to generate hydrated, vitrified samples suitable for cryo-EM. Samples were stored in liquid N_2_ to preserve the vitrified state until cryo-EM.

### Cryo-SEM and cryo-FIB-SEM

Tissues were studied in hydrated, vitrified conditions. Vitrified samples within a brass planchette were freeze-fractured with a knife in a Leica EM VCM loading station and mounted on a holder under liquid N_2_. Some samples were shuttled to a Leica ACE600 high vacuum sputter coater and etched at cryogenic conditions at <10^-6^ mbar for 15 min at −105°C. Tissues were imaged on a TESCAN S8000× Xe ion plasma focused ion beam scanning electron microscope (FIB-SEM) at 2 kV and −120°C.

Cryo-FIB-SEM allows for 3D imaging of relatively large tissue volumes at a resolution down to tens of nanometers, combining serial focused ion beam milling of vitrified samples with sequential SEM block-face imaging under cryogenic conditions^40^. After fracturing, samples were transferred to a Leica ACE600 high vacuum sputter coater and coated with a 4 nm-platinum layer to increase sample conductivity and reduce charging effects. Samples were then transferred using a Leica EM VCT500 vacuum cryo-transfer system to the FIB-SEM, where the region of interest was located using the SEM and ion beam images. The cryo-stage was tilted at 55° and stage height was adjusted to a working distance of 6 mm. Trenches of approximately 10 x 10 x 20 μm^3^ were milled and polished in front of the region of interest prior to FIB-milling and SEM image acquisition. FIB-milling parameters were 30 kV and 1 nA in the presence of flowing O_2_ applied through a gas injection system. The stack comprises 41 slices for a total volume of 10 x 10 x 4.1 μm^3^. SEM images were acquired at 2 kV, 100 pA, with pixel sizes of 5.5-19.0 nm. Unless otherwise noted, temperature was always maintained between −120 and −160 °C.

Cryo-FIB-SEM images were aligned using the “Linear Stack Alignment with SIFT” plugin in FIJI. Features of interest were isolated using manual segmentation to generate a binary mask, which was imported into MATLAB for representation as 3D smoothed isosurfaces with isotropic weighted moving average smoothing effect adjusted to match minimum voxel cube dimensions.

### Lipidomics sample preparation

Flash-frozen liver samples were homogenized in PBS at a concentration of 5 mg wet weight/mL using 1.4 mm stainless steel blend beads (Next Advance, SSB14B) in a bullet blender, taking care to avoid cross-contamination. Mass spectrometry-based lipid analysis was performed by Lipotype GmbH (Dresden, Germany) as described^80^. Lipids were extracted using a chloroform/methanol procedure^81^. Samples were spiked with internal lipid standard mixture containing: cardiolipin 14:0/14:0/14:0/14:0 (CL), ceramide 18:1;2/17:0 (Cer), diacylglycerol 17:0/17:0 (DAG), hexosylceramide 18:1;2/12:0 (HexCer), lyso-phosphatidate 17:0 (LPA), lyso-phosphatidylcholine 12:0 (LPC), lyso-phosphatidylethanolamine 17:1 (LPE), lyso-phosphatidylglycerol 17:1 (LPG), lyso-phosphatidylinositol 17:1 (LPI), lyso-phosphatidylserine 17:1 (LPS), phosphatidate 17:0/17:0 (PA), phosphatidylcholine 15:0/18:1 D7 (PC), phosphatidylethanolamine 17:0/17:0 (PE), phosphatidylglycerol 17:0/17:0 (PG), phosphatidylinositol 16:0/16:0 (PI), phosphatidylserine 17:0/17:0 (PS), cholesterol ester 16:0 D7 (CE), sphingomyelin 18:1;2/12:0;0 (SM), triacylglycerol 17:0/17:0/17:0 (TAG) and cholesterol D6 (Chol). After extraction, the organic phase was transferred to an infusion plate and dried in a speed vacuum concentrator. The dry extract was re-suspended in 7.5 mM ammonium formate in chloroform/methanol/propanol (1:2:4; V:V:V). All liquid handling steps were performed using Hamilton Robotics STARlet robotic platform with the Anti Droplet Control feature for organic solvents pipetting.

### Lipidomics MS data acquisition, analysis and post-processing

Samples were analyzed by direct infusion on a QExactive mass spectrometer (Thermo Scientific) equipped with a TriVersa NanoMate ion source (Advion Biosciences). Analysis was performed blinded. Samples were analyzed in both positive and negative ion modes with a resolution of Rm/z=200=280000 for MS and Rm/z=200=17500 for MSMS experiments, in a single acquisition. MSMS was triggered by an inclusion list encompassing corresponding MS mass ranges scanned in 1 Da increments^82^. Both MS and MSMS data were combined to monitor CE, Chol, DAG and TAG ions as ammonium adducts; LPC, LPC O-, PC and PC O-, as formiate adducts; and CL, LPS, PA, PE, PE O-, PG, PI and PS as deprotonated anions. MS only was used to monitor LPA, LPE, LPE O-, LPG and LPI as deprotonated anions, and Cer, HexCer, and SM as formiate adducts and cholesterol as ammonium adduct of an acetylated derivative^83^.

Data were analyzed with in-house developed lipid identification software based on LipidXplorer^84,85^. Data post-processing and normalization were performed using an in-house developed data management system. Only lipid identifications with a signal-to-noise ratio >5, and a signal intensity 5-fold higher than in corresponding blank samples were considered for further data analysis.

### Lipid isolation and lipid-embedded fibrin gel shear rheometry

Lipid aggregates were isolated from rat livers by adapting previously published protocols for lipid droplet isolation from rodent livers^50,51^. Briefly, rat livers were washed in PBS and minced before homogenization in 60% sucrose in TE buffer (10 mM Tris-HCl, pH 7.4; 1 mM EDTA, pH 8.0) in a Dounce homogenizer with a loose pestle. Lipid aggregates were isolated using a sucrose density gradient as described previously^50^. After isolation, the volume of lipid aggregates in solution was measured during subsequent wash steps by measuring the change in volume upon adding a known volume of dH_2_O to the isolated lipid aggregates. All steps were performed with protease and phosphatase inhibitor and on ice to prevent any degradation of lipids or associated proteins during the isolation.

Fibrin gels were prepared similarly to previously published protocols^52^. Briefly, fibrinogen isolated from bovine plasma (Sigma-Aldrich, 341573) was dissolved in 0.9% saline solution at 37 °C overnight. Thrombin isolated from bovine plasma (Sigma-Aldrich, T4648) was dissolved in 0.1% bovine serum albumin in dH_2_O, pH 6.5. To prepare fibrin gels, fibrinogen, thrombin, 200 mM HEPES solution, CaCl_2_ solution and dH_2_O were combined to yield 17 mg/mL fibrinogen, 30 mM Ca^2+^ and 2 U/mL thrombin, allowed to polymerize between the rheometer plates for 2 h at 37 °C, and then surrounded with PBS to maintain humidity. Fibrin gels (with and without lipid aggregates) were tested with a plate of 20 mm diameter and gap of 1 mm. A Peltier device incorporated into the bottom plate was used to control the sample temperature. The shear modulus was measured by applying a low oscillatory shear strain of 1% at a frequency of 10 rad/s. Axial strain was applied by changing the gap between the plates. Samples were subjected to small stepwise axial strains, between which the samples were allowed to relax completely.

### Methyl-β-cyclodextrin perfusion

Rats were anesthetized with 100 mg/kg body weight pentobarbital sodium (Sagent Pharmaceuticals, 25021-676-20). After the rat was confirmed to have reached a surgical plane of anesthesia, the abdomen was opened, the portal vein catheterized, and the inferior vena cava transected. Livers were flushed in situ with 5 mL 1000 USP units/mL heparin (Fresnius Kabi, 504015) and then with HBSS w/ Ca,Mg (Thermo Fisher, 14025) before perfusion for 2 h with HBSS w/ Ca,Mg with or without 10 mM methyl-β-cyclodextrin (Sigma-Aldrich, 332615)^53,54^. All perfusion solutions were maintained at 37 °C in a warm water bath for the duration of the perfusion. Rat liver temperature was maintained at 37 °C for the duration of the perfusion using an infrared heat lamp. After perfusion, the rat liver was removed, and samples prepared for histology and rheometry as above.

## Acknowledgements

The authors thank Xuechen Shi and Jeff Byfield for training, technical support, and feedback, and Yang Liu for assistance with the figure design. This work was carried out in part at the Center for Molecular Studies in Digestive and Liver Diseases (RRID SCR_022420), which is funded by the NIDDK under grant P30DK050306, the Cell and Developmental Biology Microscopy Core (RRID SCR_022373), and the Singh Center for Nanotechnology, which is supported by the NSF National Nanotechnology Coordinated Infrastructure Program under grant NNCI2025608.

This research was supported by funding from the NIBIB R01EB017753 (to PAJ and RGW), the American Liver Foundation Postdoctoral Research Fellowship (to DL), the NIDDK T32DK007066 (to DL), and the Center for Engineering MechanoBiology (CEMB), an NSF Science and Technology Center, under grant agreement CMMI1548571.

## References

1. Song SJ, Che-To Lai J, Lai-Hung Wong G, Wai-Sun Wong V, Cheuk-Fung Yip T. Can we use old NAFLD data under the new MASLD definition? J Hepatol. Aug 2 2023;doi:10.1016/j.jhep.2023.07.021

2. Angulo P. Nonalcoholic fatty liver disease. N Engl J Med. Apr 18 2002;346(16):1221–31. doi:10.1056/NEJMra011775

3. Younossi ZM, Stepanova M, Younossi Y, et al. Epidemiology of chronic liver diseases in the USA in the past three decades. Gut. Mar 2020;69(3):564–568. doi:10.1136/gutjnl-2019-318813

4. Chalasani N, Younossi Z, Lavine JE, et al. The diagnosis and management of nonalcoholic fatty liver disease: Practice guidance from the American Association for the Study of Liver Diseases. Hepatology. Jan 2018;67(1):328–357. doi:10.1002/hep.29367

5. Alvarez CS, Graubard BI, Thistle JE, Petrick JL, McGlynn KA. Attributable Fractions of Nonalcoholic Fatty Liver Disease for Mortality in the United States: Results From the Third National Health and Nutrition Examination Survey With 27 Years of Follow-up. Hepatology. Aug 2020;72(2):430–440. doi:10.1002/hep.31040

6. Friedman SL, Neuschwander-Tetri BA, Rinella M, Sanyal AJ. Mechanisms of NAFLD development and therapeutic strategies. Nat Med. Jul 2018;24(7):908–922. doi:10.1038/s41591-018-0104-9

7. Cotter TG, Rinella M. Nonalcoholic fatty Liver Disease 2020: The State of the Disease. Gastroenterology. May 2020;158(7):1851–1864. doi:10.1053/j.gastro.2020.01.052

8. Rinella ME, Lazarus JV, Ratziu V, et al. A multi-society Delphi consensus statement on new fatty liver disease nomenclature. J Hepatol. Jun 20 2023;doi:10.1016/j.jhep.2023.06.003

9. Perepelyuk M, Terajima M, Wang AY, et al. Hepatic stellate cells and portal fibroblasts are the major cellular sources of collagens and lysyl oxidases in normal liver and early aher injury. Am J Physiol Gastrointest Liver Physiol. Mar 15 2013;304(6):G605–14. doi:10.1152/ajpgi.00222.2012

10. Georges PC, Hui JJ, Gombos Z, et al. Increased stiffness of the rat liver precedes matrix deposition: implications for fibrosis. Am J Physiol Gastrointest Liver Physiol. Dec 2007;293(6):G1147–54. doi:10.1152/ajpgi.00032.2007

11. Olsen AL, Bloomer SA, Chan EP, et al. Hepatic stellate cells require a stiff environment for myotiibroblastic differentiation. Am J Physiol Gastrointest Liver Physiol. Jul 2011;301(1):G110–8. doi:10.1152/ajpgi.00412.2010

12. Tschumperlin DJ, Ligresti G, Hilscher MB, Shah VH. Mechanosensing and fibrosis. J Clin Invest. Jan 2 2018;128(1):74–84. doi:10.1172/JCI93561

13. Jones MG, Andriotis OG, Roberts JJ, et al. Nanoscale dysregulation of collagen structure-function disrupts mechano-homeostasis and mediates pulmonary fibrosis. Elife. Jul 3 2018;7doi:10.7554/eLife.36354

14. Kisseleva T, Brenner D. Molecular and cellular mechanisms of liver fibrosis and its regression. Nat Rev Gastroenterol Hepatol. Mar 2021;18(3):151–166. doi:10.1038/s41575-020-00372-7

15. Shoham N, Girshovitz P, Katzengold R, Shaked NT, Benayahu D, Gefen A. Adipocyte stiffness increases with accumulation of lipid droplets. Biophys J. Mar 18 2014;106(6):1421–31. doi:10.1016/j.bpj.2014.01.045

16. Yu H, Tay CY, Leong WS, Tan SC, Liao K, Tan LP. Mechanical behavior of human mesenchymal stem cells during adipogenic and osteogenic differentiation. Biochem Biophys Res Commun. Feb 26 2010;393(1):150–5. doi:10.1016/j.bbrc.2010.01.107

17. Labriola NR, Darling EM. Temporal heterogeneity in single-cell gene expression and mechanical properties during adipogenic differentiation. J Biomech. Apr 13 2015;48(6):1058–66. doi:10.1016/j.jbiomech.2015.01.033

18. Abuhattum S, Kotzbeck P, Schlussler R, et al. Adipose cells and tissues sohen with lipid accumulation while in diabetes adipose tissue stiffens. Sci Rep. Jun 20 2022;12(1):10325. doi:10.1038/s41598-022-13324-9

19. Li D, Janmey PA, Wells RG. Local fat content determines global and local stiffness in livers with simple steatosis. FASEB Bioadv. Jun 2023;5(6):251–261. doi:10.1096/la.2022-00134

20. Mueller S. Liver Elastography : Clinical Use and Interpretation. First edition 2020. ed. 2020:737.

21. Loree HM, Tobias BJ, Gibson LJ, Kamm RD, Small DM, Lee RT. Mechanical properties of model atherosclerotic lesion lipid pools. Arterioscler Thromb. Feb 1994;14(2):230–4. doi:10.1161/01.atv.14.2.230

22. Marem-Mira AC, Salomon MP, Hsu AM, Kanel GC, Golden-Mason L. Hepatic damage caused by long-term high cholesterol intake induces a dysfunctional restorative macrophage population in experimental NASH. Front Immunol. 2022;13:968366. doi:10.3389/fimmu.2022.968366

23. Tu LN, Showalter MR, Cajka T, et al. Metabolomic characteristics of cholesterol-induced non-obese nonalcoholic fatty liver disease in mice. Sci Rep. Jul 21 2017;7(1):6120. doi:10.1038/s41598-017-05040-6

24. Chiappini F, Desterke C, Bertrand-Michel J, Guemer C, Le Naour F. Hepatic and serum lipid signatures specific to nonalcoholic steatohepatitis in murine models. Sci Rep. Aug 11 2016;6:31587. doi:10.1038/srep31587

25. Horn CL, Morales AL, Savard C, Farrell GC, Ioannou GN. Role of Cholesterol-Associated Steatohepatitis in the Development of NASH. Hepatol Commun. Jan 2022;6(1):12–35. doi:10.1002/hep4.1801

26. Puri P, Baillie RA, Wiest MM, et al. A lipidomic analysis of nonalcoholic fatty liver disease. Hepatology. Oct 2007;46(4):1081–90. doi:10.1002/hep.21763

27. Min HK, Kapoor A, Fuchs M, et al. Increased hepatic synthesis and dysregulation of cholesterol metabolism is associated with the severity of nonalcoholic fatty liver disease. Cell Metab. May 2 2012;15(5):665–74. doi:10.1016/j.cmet.2012.04.004

28. Caballero F, Fernandez A, De Lacy AM, Fernandez-Checa JC, Caballeria J, Garcia-Ruiz C. Enhanced free cholesterol, SREBP-2 and StAR expression in human NASH. J Hepatol. Apr 2009;50(4):789–96. doi:10.1016/j.jhep.2008.12.016

29. Ioannou GN, Morrow OB, Connole ML, Lee SP. Association between dietary nutrient composition and the incidence of cirrhosis or liver cancer in the United States population. Hepatology. Jul 2009;50(1):175–84. doi:10.1002/hep.22941

30. Noureddin M, Zelber-Sagi S, Wilkens LR, et al. Diet Associations With Nonalcoholic fatty Liver Disease in an Ethnically Diverse Population: The Multiethnic Cohort. Hepatology. Jun 2020;71(6):1940–1952. doi:10.1002/hep.30967

31. Allard JP, Aghdassi E, Mohammed S, et al. Nutritional assessment and hepatic fatty acid composition in non-alcoholic fatty liver disease (NAFLD): a cross-sectional study. J Hepatol. Feb 2008;48(2):300–7. doi:10.1016/j.jhep.2007.09.009

32. Ichimura M, Masuzumi M, Kawase M, et al. A diet-induced Sprague-Dawley rat model of nonalcoholic steatohepatitis-related cirrhosis. J Nutr Biochem. Feb 2017;40:62–69. doi:10.1016/j.jnutbio.2016.10.007

33. Ioannou GN, Haigh WG, Thorning D, Savard C. Hepatic cholesterol crystals and crown-like structures distinguish NASH from simple steatosis. J Lipid Res. May 2013;54(5):1326–34. doi:10.1194/jlr.M034876

34. Wang L, Xu M, Jones OD, et al. Nonalcoholic fatty liver disease experiences accumulation of hepatic liquid crystal associated with increasing lipophagy. Cell Biosci. 2020;10:55. doi:10.1186/s13578-020-00414-2

35. Perepelyuk M, Chin L, Cao X, et al. Normal and Fibrotic Rat Livers Demonstrate Shear Strain Sohening and Compression stiffening: A Model for Soh Tissue Mechanics. PLoS One. 2016;11(1):e0146588. doi:10.1371/journal.pone.0146588

36. Geerligs M, Peters GW, Ackermans PA, Oomens CW, Baaijens FP. Linear viscoelastic behavior of subcutaneous adipose tissue. Biorheology. 2008;45(6):677–88.

37. Clayton EH, Garbow JR, Bayly PV. Frequency-dependent viscoelastic parameters of mouse brain tissue estimated by MR elastography. Phys Med Biol. Apr 21 2011;56(8):2391–406. doi:10.1088/0031-9155/56/8/005

38. Braun J, Tzschatzsch H, Korting C, et al. A compact 0.5 T MR elastography device and its application for studying viscoelasticity changes in biological tissues during progressive formalin fixation. Magn Reson Med. Jan 2018;79(1):470–478. doi:10.1002/mrm.26659

39. Levental I, Levental KR, Klein EA, et al. A simple indentation device for measuring micrometer-scale tissue stiffness. J Phys Condens MaJer. May 19 2010;22(19):194120. doi:10.1088/0953-8984/22/19/194120

40. Capua-Shenkar J, Varsano N, Itzhak NR, et al. Examining atherosclerotic lesions in three dimensions at the nanometer scale with cryo-FIB-SEM. Proc Natl Acad Sci U S A. Aug 23 2022;119(34):e2205475119. doi:10.1073/pnas.2205475119

41. Small DM. George Lyman Duff memorial lecture. Progression and regression of atherosclerotic lesions. Insights from lipid physical biochemistry. Arteriosclerosis. Mar-Apr 1988;8(2):103–29. doi:10.1161/01.atv.8.2.103

42. Mahamid J, Tegunov D, Maiser A, et al. Liquid-crystalline phase transitions in lipid droplets are related to cellular states and specific organelle association. Proc Natl Acad Sci U S A. Aug 20 2019;116(34):16866–16871. doi:10.1073/pnas.1903642116

43. Rogers S, Gui L, Kovalenko A, et al. Triglyceride lipolysis triggers liquid crystalline phases in lipid droplets and alters the LD proteome. J Cell Biol. Nov 7 2022;221(11)doi:10.1083/jcb.202205053

44. Lydon JE, Robinson DG. The structure of cholesteryl ester mesophases revealed by freeze-fracturing. Biochim Biophys Acta. Feb 21 1972;260(2):298–311. doi:10.1016/0005-2760(72)90041-0

45. Weibel ER, Staubli W, Gnagi HR, Hess FA. Correlated morphometric and biochemical studies on the liver cell. I. Morphometric model, stereologic methods, and normal morphometric data for rat liver. J Cell Biol. Jul 1969;42(1):68–91. doi:10.1083/jcb.42.1.68

46. Berthiaume F, Moghe PV, Toner M, Yarmush ML. Effect of extracellular matrix topology on cell structure, function, and physiological responsiveness: hepatocytes cultured in a sandwich configuration. FASEB J. Nov 1996;10(13):1471–84. doi:10.1096/fasebj.10.13.8940293

47. Duewell P, Kono H, Rayner KJ, et al. NLRP3 inflammasomes are required for atherogenesis and activated by cholesterol crystals. Nature. Apr 29 2010;464(7293):1357–61. doi:10.1038/nature08938

48. Mueller S, Millonig G, Sarovska L, et al. Increased liver stiffness in alcoholic liver disease: differentiating fibrosis from steatohepatitis. World J Gastroenterol. Feb 28 2010;16(8):966–72. doi:10.3748/wjg.v16.i8.966

49. Mueller S, Sandrin L. Liver stiffness: a novel parameter for the diagnosis of liver disease. Hepat Med. May 25 2010;2:49–67. doi:10.2147/hmer.s7394

50. Brettschneider J, Correnti JM, Lin C, et al. Rapid Lipid Droplet Isolation Protocol Using a Well-established Organelle Isolation Kit. J Vis Exp. Apr 19 2019;(146)doi:10.3791/59290

51. Ding Y, Zhang S, Yang L, et al. Isolating lipid droplets from multiple species. Nat Protoc. Jan 2013;8(1):43–51. doi:10.1038/nprot.2012.142

52. van Oosten ASG, Chen X, Chin L, et al. Emergence of tissue-like mechanics from fibrous networks confined by close-packed cells. Nature. Sep 2019;573(7772):96–101. doi:10.1038/s41586-019-1516-5

53. Rosenbaum AI, Zhang G, Warren JD, Maxfield FR. Endocytosis of beta-cyclodextrins is responsible for cholesterol reduction in Niemann-Pick type C mutant cells. Proc Natl Acad Sci U S A. Mar 23 2010;107(12):5477–82. doi:10.1073/pnas.0914309107

54. Zimmer S, Grebe A, Bakke SS, et al. Cyclodextrin promotes atherosclerosis regression via macrophage reprogramming. Sci Transl Med. Apr 6 2016;8(333):333ra50. doi:10.1126/scitranslmed.aad6100

55. Sjogren B, Hamblin MW, Svenningsson P. Cholesterol depletion reduces serotonin binding and signaling via human 5-HT(7(a)) receptors. Eur J Pharmacol. Dec 15 2006;552(1-3):1–10. doi:10.1016/j.ejphar.2006.08.069

56. Magnani F, Tate CG, Wynne S, Williams C, Haase J. Partitioning of the serotonin transporter into lipid microdomains modulates transport of serotonin. J Biol Chem. Sep 10 2004;279(37):38770–8. doi:10.1074/jbc.M400831200

57. Balberg I, Anderson CH, Alexander S, Wagner N. Excluded volume and its relation to the onset of percolation. Physical Review B. 10/01/ 1984;30(7):3933–3943. doi:10.1103/PhysRevB.30.3933

58. Balberg I, Binenbaum N, Wagner N. Percolation Thresholds in the Three-Dimensional Sticks System. Physical Review LeJers. 04/23/ 1984;52(17):1465–1468. doi:10.1103/PhysRevLett.52.1465

59. Li J, Kim JK. Percolation threshold of conducting polymer composites containing 3D randomly distributed graphite nanoplatelets. Compos Sci Technol. Aug 2007;67(10):2114–2120. doi:10.1016/j.compscitech.2006.11.010

60. Matteoni CA, Younossi ZM, Gramlich T, Boparai N, Liu YC, McCullough AJ. Nonalcoholic fatty liver disease: a spectrum of clinical and pathological severity. Gastroenterology. Jun 1999;116(6):1413–9. doi:10.1016/s0016-5085(99)70506-8

61. Wu J, Ji P, Zhang A, et al. Impact of cholesterol homeostasis within cochlear cells on auditory development and hearing loss. Front Cell Neurosci. 2023;17:1308028. doi:10.3389/fncel.2023.1308028

62. Mittal S, El-Serag HB, Sada YH, et al. Hepatocellular Carcinoma in the Absence of Cirrhosis in United States Veterans is Associated With Nonalcoholic fatty Liver Disease. Clin Gastroenterol Hepatol. Jan 2016;14(1):124–31 e1. doi:10.1016/j.cgh.2015.07.019

63. Adler M, Larocca L, Trovato FM, Marcinkowski H, Pasha Y, Taylor-Robinson SD. Evaluating the risk of hepatocellular carcinoma in patients with prominently elevated liver stiffness measurements by FibroScan: a multicentre study. HPB (Oxford). Aug 2016;18(8):678–83. doi:10.1016/j.hpb.2016.05.005

64. Abela GS, Katkoori VR, Pathak DR, et al. Cholesterol crystals induce mechanical trauma, inflammation, and neo-vascularization in solid cancers as in atherosclerosis. Am Heart J Plus. Nov 2023;35doi:10.1016/j.ahjo.2023.100317

65. Hammer SS, Dorweiler TF, McFarland D, et al. Cholesterol crystal formation is a unifying pathogenic mechanism in the development of diabetic retinopathy. Diabetologia. Sep 2023;66(9):1705–1718. doi:10.1007/s00125-023-05949-w

66. Silva GB, Gierman LM, Rakner JJ, et al. Cholesterol Crystals and NLRP3 Mediated Inflammation in the Uterine Wall Decidua in Normal and Preeclamptic Pregnancies. Front Immunol. 2020;11:564712. doi:10.3389/fimmu.2020.564712

67. Loneker AE, Alisafaei F, Kant A, et al. Lipid droplets are intracellular mechanical stressors that impair hepatocyte function. Proc Natl Acad Sci U S A. Apr 18 2023;120(16):e2216811120. doi:10.1073/pnas.2216811120

68. Shu F, Chen J, Ma X, et al. Cholesterol Crystal-Mediated Inflammation Is Driven by Plasma Membrane Destabilization. Front Immunol. 2018;9:1163. doi:10.3389/fimmu.2018.01163

69. Ichimura M, Kawase M, Masuzumi M, et al. High-fat and high-cholesterol diet rapidly induces non-alcoholic steatohepatitis with advanced fibrosis in Sprague-Dawley rats. Hepatol Res. Apr 2015;45(4):458–69. doi:10.1111/hepr.12358

70. Tan K, Cheng S, Juge L, Bilston LE. Characterising soh tissues under large amplitude oscillatory shear and combined loading. J Biomech. Apr 5 2013;46(6):1060–6. doi:10.1016/j.jbiomech.2013.01.028

71. de Jong IEM, Bodewes SB, van Leeuwen OB, et al. Restoration of Bile Duct Injury of Donor Livers During Ex Situ Normothermic Machine Perfusion. Transplantation. Feb 1 2023;doi:10.1097/TP.0000000000004531

72. Rensen SS, Slaats Y, Nijhuis J, et al. Increased hepatic myeloperoxidase activity in obese subjects with nonalcoholic steatohepatitis. Am J Pathol. Oct 2009;175(4):1473–82. doi:10.2353/ajpath.2009.080999

73. Junqueira LC, Bignolas G, Brentani RR. Picrosirius staining plus polarization microscopy, a specific method for collagen detection in tissue sections. Histochem J. Jul 1979;11(4):447–55. doi:10.1007/BF01002772

74. Furlong ST, Thibault KS, Morbelli LM, Quinn JJ, Rogers RA. Uptake and compartmentalization of fluorescent lipid analogs in larval Schistosoma mansoni. J Lipid Res. Jan 1995;36(1):1–12.

75. Ipek-Ugay S, Driessle T, Ledwig M, et al. Tabletop magnetic resonance elastography for the measurement of viscoelastic parameters of small tissue samples. J Magn Reson. Feb 2015;251:13–8. doi:10.1016/j.jmr.2014.11.009

76. de Schellenberger AA, Tzschatzsch H, Polchlopek B, et al. Sensitivity of multifrequency magnetic resonance elastography and diffusion-weighted imaging to cellular and stromal integrity of liver tissue. J Biomech. May 9 2019;88:201–208. doi:10.1016/j.jbiomech.2019.03.037

77. Kim SH, Wu SY, Baek JI, et al. A post-developmental genetic screen for zebrafish models of inherited liver disease. PLoS One. 2015;10(5):e0125980. doi:10.1371/journal.pone.0125980

78. Wisse E, Braet F, Duimel H, et al. Fixation methods for electron microscopy of human and other liver. World J Gastroenterol. Jun 21 2010;16(23):2851–66. doi:10.3748/wjg.v16.i23.2851

79. Creekmore BC, Kixmoeller K, Black BE, Lee EB, Chang YW. Ultrastructure of human brain tissue vitrified from autopsy revealed by cryo-ET with cryo-plasma FIB milling. Nat Commun. Mar 26 2024;15(1):2660. doi:10.1038/s41467-024-47066-1

80. Surma MA, Gerl MJ, Herzog R, Helppi J, Simons K, Klose C. Mouse lipidomics reveals inherent flexibility of a mammalian lipidome. Sci Rep. Sep 29 2021;11(1):19364. doi:10.1038/s41598-021-98702-5

81. Ejsing CS, Sampaio JL, Surendranath V, et al. Global analysis of the yeast lipidome by quantitative shotgun mass spectrometry. Proc Natl Acad Sci U S A. Feb 17 2009;106(7):2136–41. doi:10.1073/pnas.0811700106

82. Surma MA, Herzog R, Vasilj A, et al. An automated shotgun lipidomics plaqorm for high throughput, comprehensive, and quantitative analysis of blood plasma intact lipids. Eur J Lipid Sci Technol. Oct 2015;117(10):1540–1549. doi:10.1002/ejlt.201500145

83. Liebisch G, Binder M, Schifferer R, Langmann T, Schulz B, Schmitz G. High throughput quantification of cholesterol and cholesteryl ester by electrospray ionization tandem mass spectrometry (ESI-MS/MS). Biochim Biophys Acta. Jan 2006;1761(1):121–8. doi:10.1016/j.bbalip.2005.12.007

84. Herzog R, Schwudke D, Schuhmann K, et al. A novel informatics concept for high-throughput shotgun lipidomics based on the molecular fragmentation query language. Genome Biol. 2011;12(1):R8. doi:10.1186/gb-2011-12-1-r8

85. Herzog R, Schuhmann K, Schwudke D, et al. LipidXplorer: a sohware for consensual cross-plaqorm lipidomics. PLoS One. 2012;7(1):e29851. doi:10.1371/journal.pone.0029851

